# BAP1 constrains pervasive H2AK119ub1 to control the transcriptional potential of the genome

**DOI:** 10.1101/2020.11.13.381251

**Authors:** Nadezda A. Fursova, Anne H. Turberfield, Neil P. Blackledge, Emma L. Findlater, Anna Lastuvkova, Miles K. Huseyin, Paula Dobrinić, Robert J. Klose

**Affiliations:** Department of Biochemistry, University of Oxford, South Parks Rd, Oxford, OX1 3QU, United Kingdom

## Abstract

Histone-modifying systems play fundamental roles in gene regulation and the development of multicellular organisms. Histone modifications that are enriched at gene regulatory elements have been heavily studied, but the function of modifications that are found more broadly throughout the genome remains poorly understood. This is exemplified by histone H2A mono-ubiquitylation (H2AK119ub1) which is enriched at Polycomb-repressed gene promoters, but also covers the genome at lower levels. Here, using inducible genetic perturbations and quantitative genomics, we discover that the BAP1 deubiquitylase plays an essential role in constraining H2AK119ub1 throughout the genome. Removal of BAP1 leads to pervasive accumulation of H2AK119ub1, which causes widespread reductions in gene expression. We show that elevated H2AK119ub1 represses gene expression by counteracting transcription initiation from gene regulatory elements, causing reductions in transcription-associated histone modifications. Furthermore, failure to constrain pervasive H2AK119ub1 compromises Polycomb complex occupancy at a subset of Polycomb target genes leading to their derepression, therefore explaining the original genetic characterisation of BAP1 as a Polycomb group gene. Together, these observations reveal that the transcriptional potential of the genome can be modulated by regulating the levels of a pervasive histone modification, without the need for elaborate gene-specific targeting mechanisms.

## Introduction

In eukaryotes, DNA is wrapped around histones to form nucleosomes and chromatin which packages the genome inside the nucleus. In addition to their structural role, histones are subject to a variety of post-translational modifications (PTMs), which have been proposed to play important roles in regulation of gene expression and other chromosomal processes (Kouzarides 2007; Bannister and Kouzarides 2011; Groth et al. 2007; Zhao and Garcia 2015; Hauer and Gasser 2017). If chromatin-modifying systems are perturbed, this can lead to profound alterations in gene expression resulting in severe developmental disorders and cancer (Audia and Campbell 2016; Atlasi and Stunnenberg 2017; Zhao and Shilatifard 2019; Bracken et al. 2019). However, for many histone modifications, the mechanisms that control their levels throughout the genome and ultimately how this influences gene expression remains poorly understood.

Genome-wide profiling has revealed that some histone modifications are specifically enriched at gene promoters and distal regulatory elements (Barski et al. 2007; Ernst and Kellis 2010; Kharchenko et al. 2011; Zhou et al. 2011; Ho et al. 2014), where they have been proposed to regulate chromatin accessibility and work with the transcriptional machinery to control gene expression (Lee et al. 1993; Vettese-Dadey et al. 1996; Pray-Grant et al. 2005; Wysocka et al. 2006; Vermeulen et al. 2007; Lauberth et al. 2013; Zhang et al. 2017a). However, it has also emerged that there are other histone modifications that are extremely abundant and cover broad regions of the genome, extending far beyond genes and gene regulatory elements (Kharchenko et al. 2011; Ferrari et al. 2014; Lee et al. 2015; Kahn et al. 2016; Zheng et al. 2016; Carelli et al. 2017; Fursova et al. 2019). Much less effort has been placed on studying these more pervasive histone modifications, raising the possibility that they could also have important and previously underappreciated roles in gene regulation.

The Polycomb Repressive Complex 1 (PRC1) is an E3 ubiquitin ligase that catalyzes mono-ubiquitylation of histone H2A (H2AK119ub1) (Wang et al. 2004; de Napoles et al. 2004; Buchwald et al. 2006). PRC1 is targeted to CpG island-associated gene promoters where it can deposit high levels of H2AK119ub1 (Ku et al. 2008; Farcas et al. 2012; He et al. 2013; Wu et al. 2013; Bauer et al. 2016), and this is central to Polycomb-mediated gene repression (Endoh et al. 2012; Blackledge et al. 2014; Tsuboi et al. 2018; Fursova et al. 2019; Blackledge et al. 2020; Tamburri et al. 2020). A second Polycomb Repressive Complex, PRC2, is recruited to the same sites (Boyer et al. 2006; Bracken 2006; Li et al. 2017a; Perino et al. 2018) where it deposits histone H3 lysine 27 methylation (H3K27me3) (Cao et al. 2002; Kuzmichev 2002; Czermin et al. 2002; Müller et al. 2002), leading to the formation of transcriptionally repressive Polycomb chromatin domains that have high levels of PRC1, PRC2, and their respective histone modifications (Mikkelsen et al. 2007; Ku et al. 2008). In addition to this punctate high-level enrichment of H2AK119ub1 at Polycomb target gene promoters, we and others have recently demonstrated that H2AK119ub1 is also found broadly throughout the genome, albeit at much lower levels (Lee et al. 2015; Kahn et al. 2016; Fursova et al. 2019). However, whether this genome-wide pool of H2AK119ub1 influences gene expression has remained enigmatic.

Interestingly, H2AK119ub1 is highly dynamic (Seale 1981) and a number of deubiquitylating enzymes (DUBs) have been proposed to regulate its levels (reviewed in Belle and Nijnik 2014; Aquila and Atanassov 2020). The most extensively characterised and evolutionary conserved of these DUBs is BAP1, which interacts with ASXL proteins to form the Polycomb Repressive Deubiquitinase complex (PR-DUB) (Scheuermann et al. 2010; Wu et al. 2015; Sahtoe et al. 2016; Kloet et al. 2016; Hauri et al. 2016; Campagne et al. 2019). Previous attempts to understand how BAP1 regulates gene expression and whether this relies on its H2AK119ub1 deubiquitylase activity have primarily focused on examining how the PR-DUB complex is targeted to gene promoters and distal regulatory elements, and how this regulates binding and/or activity of chromatin-modifying transcriptional co-activators (Li et al. 2017b; Wang et al. 2018; Campagne et al. 2019; Kuznetsov et al. 2019; Kolovos et al. 2020; Szczepanski et al. 2020). While this has revealed that BAP1 can remove H2AK119ub1 at specific loci, its primary site of action in the genome and the mechanisms by which it controls gene expression have appeared to be context-dependent and in some cases difficult to reconcile with the known roles of H2AK119ub1 in gene regulation. Therefore, how H2AK119ub1 levels in the genome are modulated by BAP1 and how this influences transcription remains poorly defined. Addressing these questions is particularly important in the light of the essential role that BAP1 plays as a tumour suppressor (Ventii et al. 2008; Dey et al. 2012; Carbone et al. 2013; Murali et al. 2013; Daou et al. 2015), and could provide important new insight into how BAP1 dysfunction causes cellular transformation.

To dissect how BAP1 controls H2AK119ub1 levels and gene expression, here we integrate genome editing, inducible genetic perturbations, and quantitative genomics. We discover that BAP1 functions to constrain pervasive H2AK119ub1 throughout the genome, with no preference for gene promoters or distal regulatory elements. We demonstrate that by counteracting pervasive H2AK119ub1, BAP1 plays a fundamental role in facilitating gene expression. In the absence of BAP1, elevated H2AK119ub1 broadly inhibits transcription initiation from gene regulatory elements and causes widespread reductions in transcription-associated histone modifications without limiting chromatin accessibility. Finally, we discover that a subset of Polycomb target genes rely on BAP1 for their silencing and provide a mechanistic rationale for how BAP1 can indirectly support Polycomb-mediated gene repression. Together, these observations demonstrate how the levels of a pervasive histone modification must be appropriately controlled to enable the transcriptional potential of the genome.

## Results

### BAP1 functions pervasively throughout the genome to constrain H2AK119ub1

Given our recent discovery that H2AK119ub1 is deposited more broadly throughout the genome than previously appreciated (Fursova et al. 2019), we set out to determine where in the genome BAP1 functions to control the levels of H2AK119ub1 and how this influences gene expression. To address these important questions, we developed a BAP1 conditional knockout mouse embryonic stem cell (ESC) line (*Bap1*^*fl/fl*^) in which addition of tamoxifen (OHT) enables inducible removal of BAP1, allowing us to capture the primary effects that BAP1 loss has on H2AK119ub1 and gene expression. Importantly, tamoxifen treatment of *Bap1*^*fl/fl*^ cells resulted in a complete loss of BAP1 protein, while the levels of BAP1-interacting partners were largely unchanged (Fig. 1A). In line with previous observations in BAP1 knockout mouse ESCs and human cancer cell lines (Wang et al. 2018; He et al. 2019; Campagne et al. 2019), western blot analysis showed that H2AK119ub1 levels were markedly increased (~ 50%) following BAP1 removal, whereas H2BK120ub1 was unaffected (Fig. 1B).

**Figure 1.**
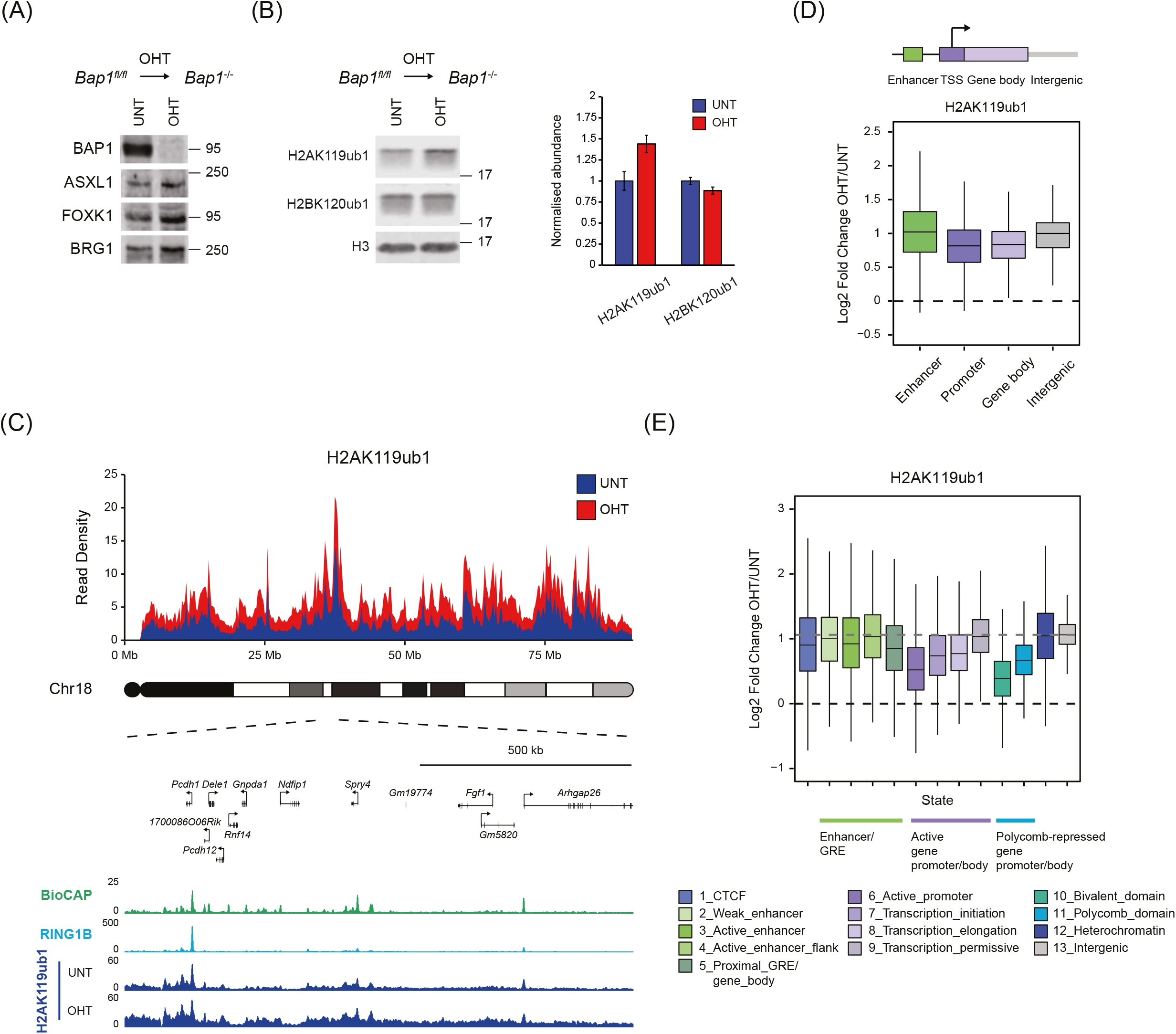
BAP1 functions pervasively throughout the genome to constrain H2AK119ub1. **(A)** Western blot analysis for BAP1 and other subunits of the PR-DUB complex (ASXL1 and FOXK1) in untreated (UNT) and OHT-treated (OHT) *Bap1*^*fl/fl*^ ESCs. BRG1 is shown as a loading control. **(B)** Western blot analysis (*left panel*) and quantification (*right panel*) of H2AK119ub1 and H2BK120ub1 levels relative to histone H3 in untreated (UNT) and OHT-treated (OHT) *Bap1*^*fl/fl*^ ESCs. Error bars represent SEM (n = 3). **(C)** A chromosome density plot showing H2AK119ub1 cChIP-seq signal across chromosome 18 in *Bap1*^*fl/fl*^ ESCs (untreated and OHT-treated) with an expanded snapshot of a region on chromosome 18 shown below. BioCAP-seq and RING1B cChIP-seq in wild-type ESCs are also shown to indicate the location of CGIs that are occupied by PRC1. **(D)** Boxplots comparing log2-fold changes in H2AK119ub1 cChIP-seq signal at gene regulatory elements (enhancers and promoters), gene bodies, and intergenic regions in *Bap1*^*fl/fl*^ ESCs following OHT treatment. **(E)** Boxplots comparing log2-fold changes in H2AK119ub1 cChIP-seq signal following OHT treatment in *Bap1*^*fl/fl*^ ESCs across different chromatin states derived from unsupervised genome segmentation using ChromHMM. Chromatin states are grouped based on the underlying gene regulatory elements (GREs) and transcriptional activity. The dashed grey line indicates the overall change in H2AK119ub1 levels in the genome as determined by its median value in intergenic regions.

Having shown that conditional knockout of BAP1 leads to an increase in H2AK119ub1 (Fig. 1B), we set out to define where in the genome H2AK119ub1 was elevated using an unbiased quantitative genomic approach. To achieve this, we carried out calibrated ChIP-seq (cChIP-seq) for H2AK119ub1 before and after removal of BAP1. Remarkably, this revealed a major and widespread accumulation of H2AK119ub1, which was evident when we visualised changes in H2AK119ub1 across an entire chromosome and also when we focused on individual regions of chromosomes (Fig. 1C; Supplemental Fig. S1D). Importantly, the magnitude of H2AK119ub1 accumulation appeared to be largely uniform throughout the genome (Fig. 1C), showing no preference for gene regulatory elements, including promoters and enhancers (Fig. 1D; Supplemental Fig. S1A), where BAP1 has been previously proposed to function (Wang et al. 2018; Kuznetsov et al. 2019; Campagne et al. 2019). To characterise the effect of BAP1 removal on H2AK119ub1 in more detail, we employed an unsupervised ChromHMM classification approach (Ernst and Kellis 2012) to segment the genome into 13 chromatin states encompassing all major functional genomic annotations (Supplemental Fig. S1B) and examined changes in H2AK119ub1 across these distinct states. This revealed that while the effects on H2AK119ub1 were slightly less pronounced at Polycomb-enriched genomic regions and those encompassing actively transcribed genes, all chromatin states were significantly affected and gained similar amounts of H2AK119ub1 (Fig. 1E; Supplemental Fig. S1C,D). Together, these observations demonstrate that BAP1 functions pervasively and indiscriminately throughout the genome to constrain H2AK119ub1.

### Pervasive accumulation of H2AK119ub1 in the absence of BAP1 causes widespread reductions in gene expression

Given that H2AK119ub1 plays a central role in PRC1-mediated gene repression (Endoh et al. 2012; Blackledge et al. 2020; Tamburri et al. 2020), we were curious to determine what effect BAP1 removal and the resulting accumulation of H2AK119ub1 throughout the genome would have on gene expression. Therefore, we carried out calibrated nuclear RNA sequencing (cnRNA-seq) in our conditional BAP1 knockout cells. This revealed that removal of BAP1 caused surprisingly widespread changes in gene expression, with the majority of genes exhibiting reduced expression (Fig. 2A). Remarkably, we found that approximately 6500 genes showed at least a 20% reduction in expression, of which 2828 genes were significantly reduced by at least 1.5-fold, indicating that BAP1 plays a broad role in promoting gene expression.

**Figure 2.**
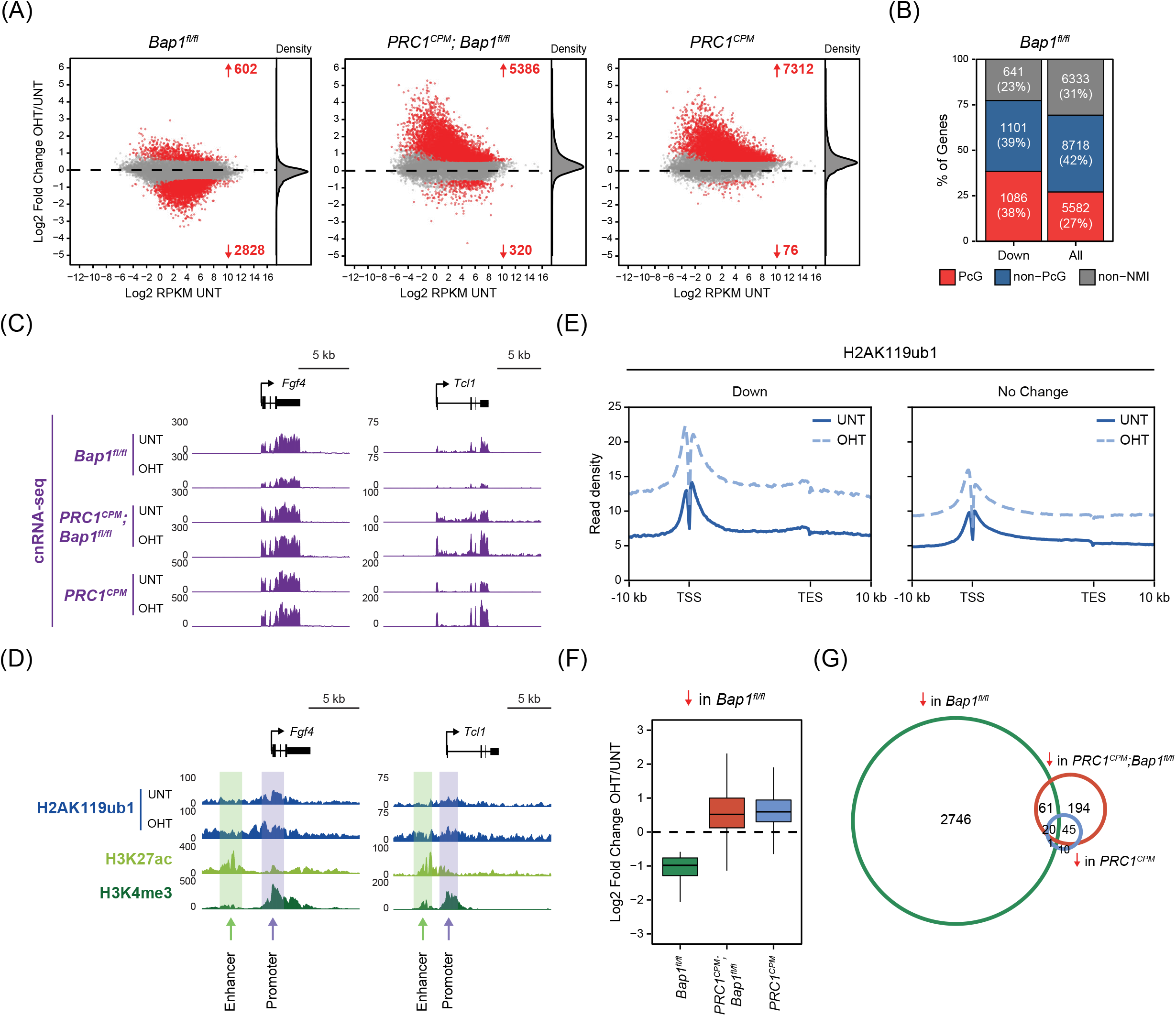
Pervasive accumulation of H2AK119ub1 in the absence of BAP1 causes widespread reductions in gene expression. **(A)** MA-plots showing log2-fold changes in gene expression (cnRNA-seq) in *Bap1*^*fl/fl*^, *PRC1*^*CPM*^;*Bap1*^*fl/fl*^ and *PRC1*^*CPM*^ ESCs following OHT treatment. Significant gene expression changes (p-adj < 0.05 and > 1.5-fold) are shown in red. The density of gene expression changes is shown on the right. **(B)** A bar plot illustrating the distribution of different gene classes among genes showing significantly reduced expression following OHT treatment in *Bap1*^*fl/fl*^ ESCs based on cnRNA-seq analysis (p-adj < 0.05 and > 1.5-fold). PcG corresponds to Polycomb-occupied genes; Non-PcG to non-Polycomb-occupied genes; Non-NMI to genes lacking a non-methylated CGI (NMI) at their promoter. **(C)** Snapshots of genes whose expression is significantly reduced (p-adj < 0.05 and > 1.5-fold) following removal of BAP1, showing gene expression (cnRNA-seq) in *Bap1*^*fl/fl*^, *PRC1*^*CPM*^;*Bap1*^*fl/fl*^ and *PRC1*^*CPM*^ ESCs (untreated and OHT-treated). **(D)** Snapshots of genes whose expression is significantly reduced (p-adj < 0.05 and > 1.5-fold) following removal of BAP1, showing H2AK119ub1 cChIP-seq in *Bap1*^*fl/fl*^ ESCs (untreated and OHT-treated). Also shown is cChIP-seq for H3K27ac and H3K4me3 in untreated *Bap1*^*fl/fl*^ ESCs to highlight the position of promoters (H3K27ac-high, H3K4me3-high) and nearest putative enhancers (H3K27ac-high, H3K4me3-low) for these genes. **(E)** Metaplots of H2AK119ub1 cChIP-seq signal in *Bap1*^*fl/fl*^ ESCs (untreated and OHT-treated) across genes that show a significant reduction (Down, n = 2828) or no change (No Change, n = 17,203) in expression following BAP1 removal based on cnRNA-seq analysis (p-adj < 0.05 and > 1.5-fold). **(F)** Boxplots comparing log2-fold changes in expression (cnRNA-seq) following OHT treatment in *Bap1*^*fl/fl*^ *(green), PRC1*^*CPM*^;*Bap1*^*fl/fl*^ *(red)* and *PRC1*^*CPM*^ *(blue)* ESCs for genes whose expression is significantly reduced (p-adj < 0.05 and > 1.5-fold) in the absence of BAP1. **(G)** A Venn diagram showing the overlap between genes that show a significant reduction in expression based on cnRNA-seq analysis (p-adj < 0.05 and > 1.5-fold) following OHT treatment in *Bap1*^*fl/fl*^ *(green), PRC1*^*CPM*^;*Bap1*^*fl/fl*^ *(red)* and *PRC1*^*CPM*^ *(blue)* ESCs.

Although reductions in gene expression following BAP1 removal were widespread, expression of some genes was more severely affected than others. Importantly, the majority of genes showing significantly reduced expression were not classical Polycomb target genes (Fig. 2B; Supplemental Fig. S2A). However, interestingly, these genes were often found in regions of the genome which had higher levels of H2AK119ub1 in wild-type cells, and in the absence of BAP1 also acquired higher levels of H2AK119ub1 than genes that were not significantly affected (Fig. 2C-E; Supplemental Fig. S2B-E). Importantly, the increase in H2AK119ub1 was not specific to the promoters or enhancers of these genes, but was evident across the entire gene and flanking regions (Fig. 2D,E; Supplemental Fig. S2B-E). Together, these observations suggest that widespread reductions in gene expression following BAP1 removal likely result from pervasive accumulation of H2AK119ub1, with some genes being more susceptible to these effects than others.

To directly test whether elevated H2AK119ub1 was responsible for gene repression in the absence of BAP1, we developed an inducible mouse ES cell line (*PRC1*^*CPM*^;*Bap1*^*fl/fl*^) in which we could simultaneously disrupt BAP1 and inactivate PRC1 catalysis to remove H2AK119ub1 (Supplemental Fig. S2F,G) (Blackledge et al. 2020). We then carried out cnRNA-seq and compared the effects on gene expression caused by concurrent removal of BAP1 and H2AK119ub1 with the effects caused by removing BAP1 or H2AK119ub1 individually (Fig. 2A; Supplemental Fig. S2H). Strikingly, in the absence of H2AK119ub1, removal of BAP1 no longer caused widespread reductions in gene expression (Fig. 2A,C,F,G; Supplemental Fig. S2I), indicating that H2AK119ub1 was required for these effects. In contrast, Polycomb target genes were derepressed following catalytic inactivation of PRC1 regardless of whether BAP1 was disrupted (Fig. 2A; Supplemental Fig. S2J). Therefore, we conclude that BAP1 counteracts accumulation of H2AK119ub1 throughout the genome, and in its absence elevated H2AK119ub1 causes widespread inhibition of gene expression.

### BAP1 counteracts pervasive H2AK119ub1 to promote transcription initiation from gene regulatory elements

To understand how accumulation of H2AK119ub1 counteracts gene expression, we examined how the function of RNA Polymerase II (Pol II) was affected after BAP1 removal. To achieve this, we carried out cChIP-seq to quantitate total Pol II levels and also examined its phosphorylation states which are associated with either transcription initiation (Ser5P) or elongation (Ser2P) (Buratowski 2009; Harlen and Churchman 2017). When we inspected genes whose expression was significantly reduced following BAP1 removal, we found that levels of Pol II and its phosphorylated forms were decreased at promoters and over gene bodies (Fig. 3A; Supplemental Fig. S3A,B). The reduction in Ser2P in gene bodies was similar in magnitude to the decrease in Pol II levels (Fig. 3A; Supplemental Fig. S3A,D), indicating that elongation-associated phosphorylation was not specifically disrupted, despite reduced transcription. In contrast, the reduction in Ser5P at the promoters of these genes was larger in magnitude than the decrease in Pol II occupancy (Fig. 3A; Supplemental Fig. S3A,D), suggesting that elevated H2AK119ub1 limits transcription initiation which leads to reduced transcription and gene expression.

**Figure 3.**
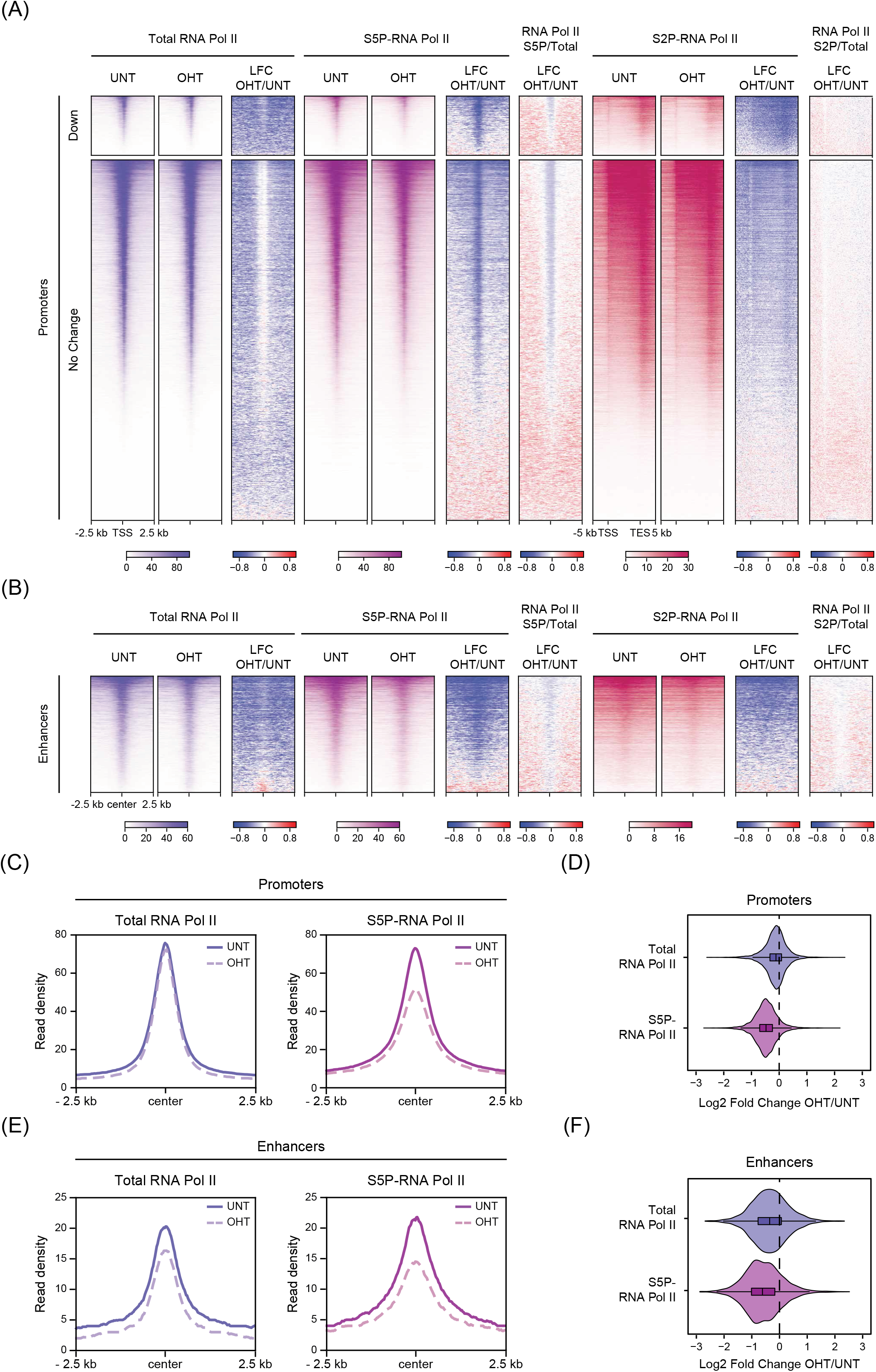
BAP1 counteracts pervasive H2AK119ub1 to promote transcription initiation from gene regulatory elements. **(A)** Heatmaps illustrating cChIP-seq signal for total Pol II occupancy and its Ser5-phosphorylation (S5P) at gene promoters, as well as Pol II Ser2-phosphorylation (S2P) over gene bodies in *Bap1*^*fl/fl*^ ESCs (untreated and OHT-treated). Also shown are the log2-fold changes in cChIP-seq signal after BAP1 removal (LFC OHT/UNT) and the log2-fold changes in the abundance of Ser5- and Ser2-phosphorylation relative to total Pol II levels (S5P/Total and S2P/Total). Genes were segregated into those that show a significant reduction (Down, n = 2828) or no change (No Change, n = 17,203) in expression following BAP1 removal based on cnRNA-seq analysis. Intervals were sorted by total Pol II cChIP-seq signal in untreated *Bap1*^*fl/fl*^ ESCs. **(B)** Heatmaps illustrating cChIP-seq signal for total Pol II occupancy, as well as its Ser5P and Ser2P forms, at active enhancers in *Bap1*^*fl/fl*^ ESCs (untreated and OHT-treated). As in (A), the log2-fold changes in cChIP-seq signal after BAP1 removal (LFC OHT/UNT) are shown together with the log2-fold changes in the abundance of Ser5P and Ser2P relative to total Pol II levels (S5P/Total and S2P/Total). Intervals were sorted by total Pol II cChIP-seq signal in untreated *Bap1*^*fl/fl*^ ESCs. **(C)** Metaplots of total and Ser5-phosphorylated Pol II cChIP-seq signal at active gene promoters in *Bap1*^*fl/fl*^ ESCs (untreated and OHT-treated). **(D)** Violinplots comparing log2-fold changes in total and Ser5-phosphorylated Pol II cChIP-seq signal at active gene promoters in *Bap1*^*fl/fl*^ ESCs following OHT treatment. **(E)** As in (C) but for active enhancers. **(F)** As in (D) but for active enhancers.

Given that removal of BAP1 caused pervasive accumulation of H2AK119ub1 throughout the genome (Fig. 1), we wondered whether the repressive effects of this histone modification on transcription may in fact extend beyond the subset of genes that showed significant reductions in gene expression. When we examined genes whose expression did not change significantly after BAP1 removal, we found that the occupancy of Pol II at their promoters was only modestly affected, but there were widespread reductions in the levels of Pol II and Ser2P over gene bodies (Fig. 3A; Supplemental Fig. S3A,B), which were similar in magnitude (Fig. 3A; Supplemental Fig. S3D). Importantly, these effects on Pol II in the gene body correlated well with changes in transcript levels (Supplemental Fig. S3C), indicating that elevated H2AK119ub1 causes widespread reductions in transcription and gene expression (Supplemental Fig. S3E), despite only a subset of genes being captured as having significantly reduced expression in cnRNA-seq analysis. Importantly, in contrast to Pol II occupancy which was only modestly affected, transcription initiation-associated Ser5P was markedly reduced at the promoters of all genes, including those whose expression did not change significantly after BAP1 removal (Fig. 3A,C,D; Supplemental Fig. S3A,B,F). This indicates that the widespread reductions in transcription following BAP1 removal likely result from a reduced capacity of Pol II to initiate transcription. Since the accumulation of H2AK119ub1 in the absence of BAP1 is not restricted to genes or their promoters (Fig. 1D), we wondered whether the observed effects on Pol II function may in fact extend to other gene regulatory elements, like enhancers, which have been also reported to bind Pol II and initiate transcription (Li et al. 2016; Andersson and Sandelin 2020; Sartorelli and Lauberth 2020). This revealed that there was a pronounced decrease in total Pol II occupancy and an even larger reduction in Ser5P at enhancers (Fig. 3B,E,F; Supplemental Fig. S3A,F). Together, these observations demonstrate that BAP1 functions broadly throughout the genome to support transcription initiation from gene regulatory elements by constraining pervasive H2AK119ub1.

### Aberrant accumulation of H2AK119ub1 compromises transcription-associated histone modifications but does not limit chromatin accessibility

Having established that elevated H2AK119ub1 in the absence of BAP1 broadly inhibits transcription initiation from promoters and enhancers (Fig. 3), we wanted to investigate whether chromatin features associated with transcription were also affected. To address this question, we carried out cChIP-seq for histone modifications that are typically enriched at active promoters (H3K27ac and H3K4me3) or enhancers (H3K27ac and H3K4me1) (Calo and Wysocka 2013; Andersson and Sandelin 2020). Interestingly, we observed a widespread decrease in H3K27ac at both types of gene regulatory elements in the absence of BAP1, with enhancers showing more pronounced reductions (Fig. 4A-D; Supplemental Fig. S4A-D). Removal of BAP1 also compromised H3K4me3 at gene promoters, but this effect was on average much more modest and mostly limited to genes that showed significant reductions in expression (Fig. 4A,C; Supplemental Fig. S4A,B,D). In contrast, H3K4me3 at enhancers was markedly reduced, despite the starting levels of this modification being considerably lower than at promoters (Fig. 4A,B,D; Supplemental Fig. S4A-C). Finally, we also observed a modest but widespread decrease in H3K4me1 around promoters and enhancers, which was accompanied by a slight increase at the center of these regulatory elements (Fig. 4A-D; Supplemental Fig. S4A-D). Therefore, we find that BAP1 removal leads to moderate and seemingly indiscriminate effects on transcription-associated histone modifications at both promoters and enhancers, which correlate with the effects on gene expression (Supplemental Fig. S4E). This observation differs from previous studies that have implicated BAP1 and other PR-DUB subunits in directly recruiting chromatin-modifying transcriptional co-activators to either promoters or enhancers to specifically affect histone modifications at these sites (Li et al. 2017b; Wang et al. 2018; Szczepanski et al. 2020). Instead, our new findings are more consistent with a model in which elevated H2AK119ub1 inhibits transcription initiation in the absence of BAP1, which then leads to modest but broad effects on transcription-associated histone modifications.

**Figure 4.**
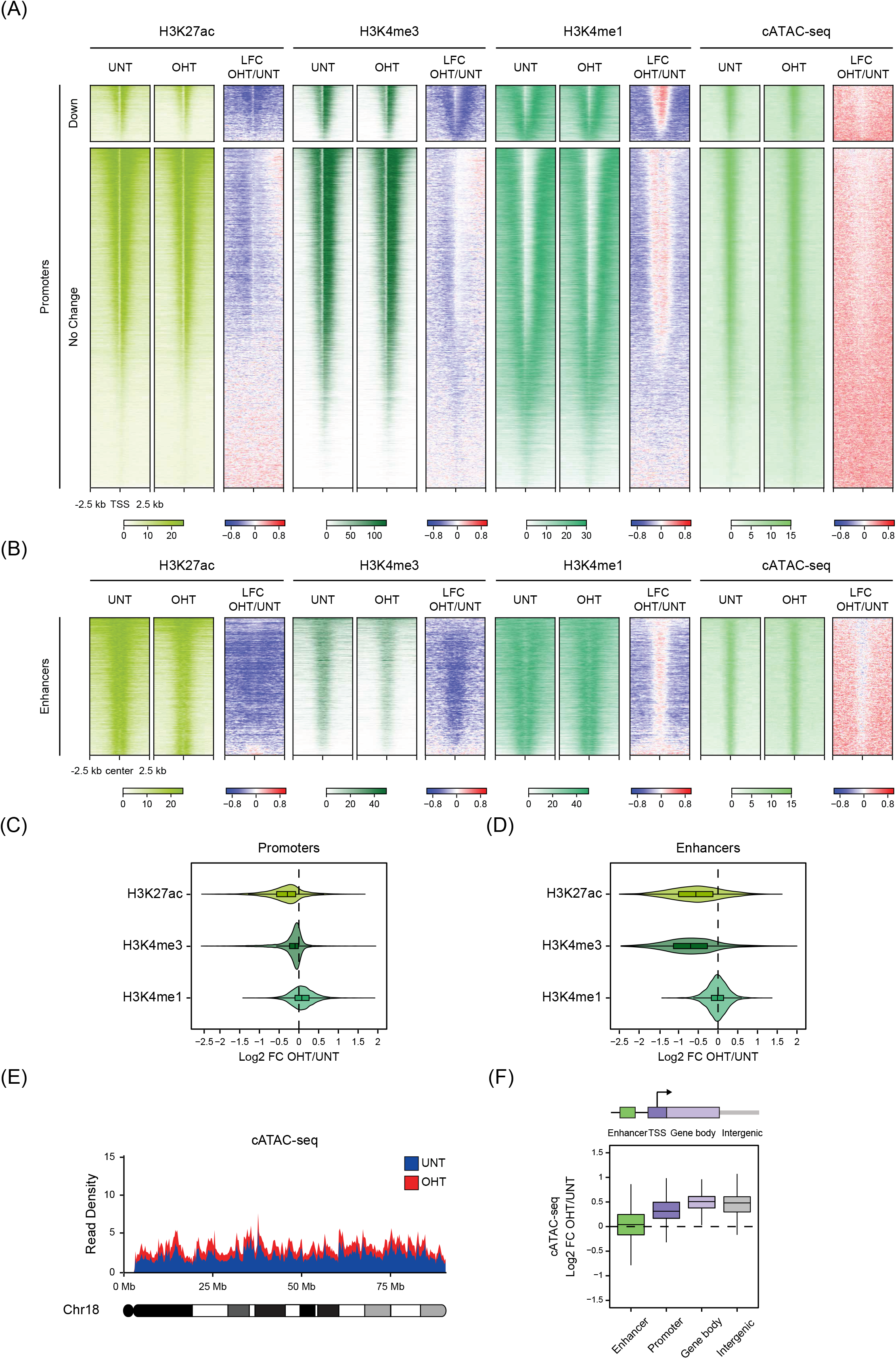
Aberrant accumulation of H2AK119ub1 compromises transcription-associated histone modifications but not chromatin accessibility at gene regulatory elements. **(A)** Heatmaps illustrating H3K27ac, H3K4me3, and H3K4me1 cChIP-seq signal at gene promoters in *Bap1*^*fl/fl*^ ESCs (untreated and OHT-treated). cATAC-seq is shown as a measure of chromatin accessibility. Also shown are the log2-fold changes in cChIP-seq and cATAC-seq signal after BAP1 removal (LFC OHT/UNT). Genes were segregated into those that show a significant reduction (Down, n = 2828) or no change (No Change, n = 17,203) in expression following BAP1 removal based on cnRNA-seq analysis. Intervals were sorted by total Pol II cChIP-seq signal in untreated *Bap1*^*fl/fl*^ ESCs. **(B)** As in (A) but for active enhancers. **(C)** Violinplots comparing log2-fold changes in cChIP-seq signal for H3K27ac, H3K4me3 and H3K4me1 at active gene promoters in *Bap1*^*fl/fl*^ ESCs following OHT treatment. **(D)** As in (C) but for active enhancers. **(E)** A chromosome density plot showing chromatin accessibility across chromosome 18 as measured by cATAC-seq in *Bap1*^*fl/fl*^ ESCs (untreated and OHT-treated). This illustrates a widespread increase in cATAC-seq signal throughout the genome following BAP1 removal. **(F)** Boxplots comparing log2-fold changes in cATAC-seq signal at gene regulatory elements (enhancers and promoters), gene bodies, and intergenic regions in *Bap1*^*fl/fl*^ ESCs following OHT treatment.

Given that some chromatin modifications have been proposed to function through making chromatin less accessible to gene regulatory factors (Francis et al. 2001, 2004; Danzer and Wallrath 2004; Soufi et al. 2012; Fyodorov et al. 2018; Becker et al. 2016), we sought to determine whether elevated H2AK119ub1 could elicit its widespread effects on transcription by limiting chromatin accessibility. To test this possibility, we performed calibrated ATAC-seq (cATAC-seq) that measures chromatin accessibility by its susceptibility to tagmentation by Tn5 transposase (Buenrostro et al. 2015). Importantly, we found that accumulation of H2AK119ub1 in the absence of BAP1 did not cause major reductions in chromatin accessibility at gene promoters and enhancers (Fig. 4A,B,F; Supplemental Fig. S4A-D). Instead, to our surprise, we found that following BAP1 removal chromatin accessibility was modestly increased throughout the genome, in a similar manner to the pervasive accumulation of H2AK119ub1 (Fig. 4A,B,E,F). Importantly, this demonstrates that H2AK119ub1 does not counteract transcription simply by limiting the access of regulatory factors to promoters and enhancers. Instead, pervasive accumulation of H2AK119ub1 in the absence of BAP1 leads to widespread reductions in transcription initiation and its associated histone modifications. Together, our findings illustrate how a pervasive histone modification that can inhibit transcription initiation must be appropriately controlled to support the transcriptional potential of the genome.

### BAP1 indirectly supports repression of a subset of Polycomb target genes

Our discovery that BAP1 constrains pervasive H2AK119ub1 to facilitate gene expression is conceptually at odds with genetic characterisation of the *Drosophila* orthologues of BAP1 (Calypso) and other PR-DUB components as Polycomb group (PcG) transcriptional repressors (Jürgens 1985; Soto et al. 1995; de Ayala Alonso et al. 2007; Scheuermann et al. 2010). Intriguingly, despite the majority of genes showing reduced expression in BAP1-deficient cells, we also identified 602 genes whose expression was significantly increased in the absence of BAP1 (Fig. 2A). Remarkably, when we examined these genes in more detail, we found that the majority were Polycomb target genes, enriched in GO categories related to regulation of developmental processes which are characteristic of PRC1-repressed genes in mouse ESCs (Fig. 5A-D). Therefore, we show that BAP1 is required to repress a subset of Polycomb target genes, consistent with its genetic designation as a PcG gene.

**Figure 5.**
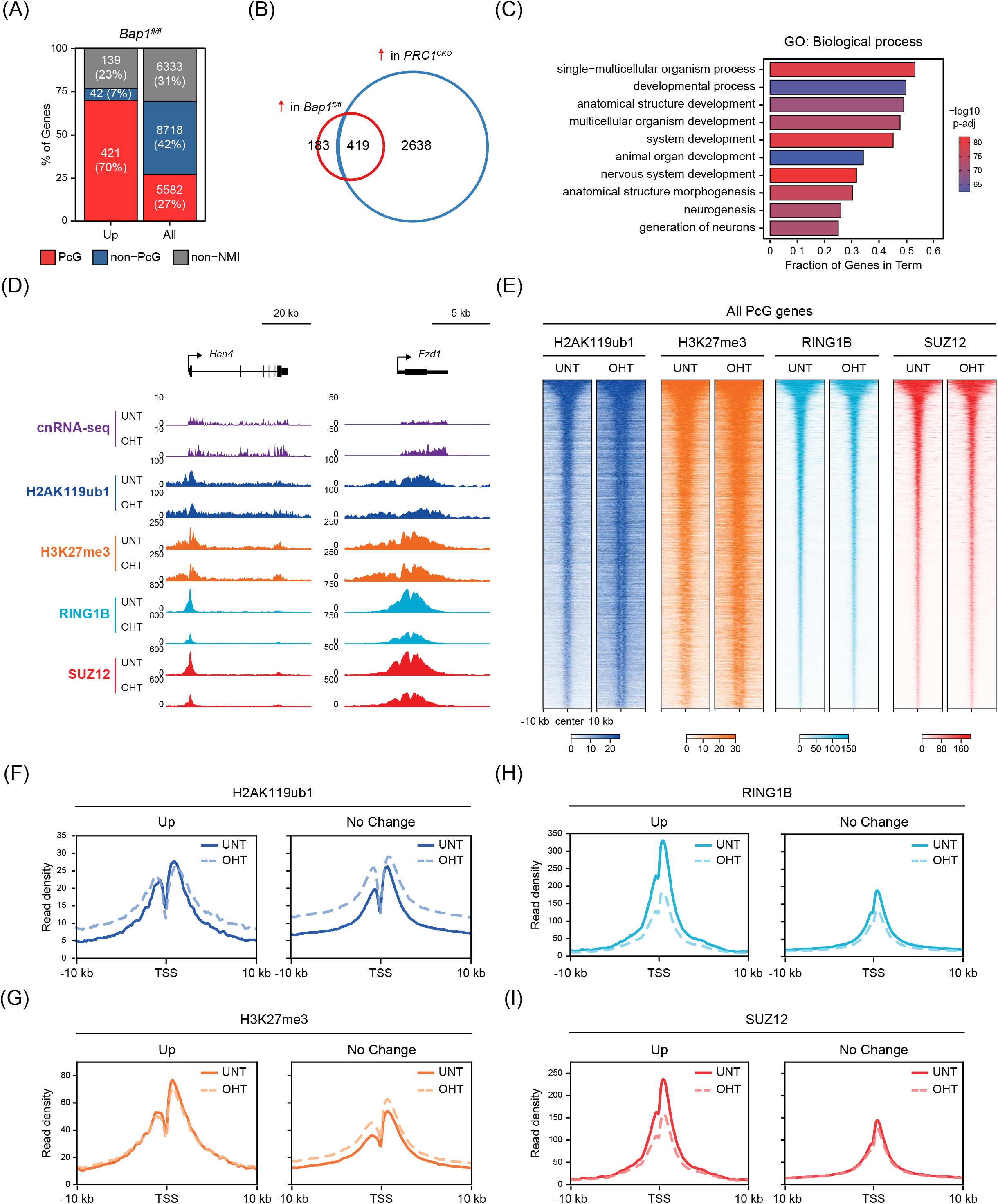
BAP1 indirectly supports repression of a subset of Polycomb target genes by counteracting pervasive H2AK119ub1 to focus Polycomb complex occupancy at target sites. **(A)** A bar plot illustrating the distribution of different gene classes among genes that become significantly derepressed (p-adj < 0.05 and > 1.5-fold) following OHT treatment in *Bap1*^*fl/fl*^ ESCs. PcG corresponds to Polycomb-occupied genes; Non-PcG to non-Polycomb-occupied genes; Non-NMI to genes lacking a non-methylated CGI (NMI) at their promoter. **(B)** A Venn diagram showing the overlap between genes that become significantly derepressed (p-adj < 0.05 and > 1.5-fold) following OHT treatment in *Bap1*^*fl/fl*^ *(red)* or *PRC1*^*CKO*^ *(blue)* ESCs. **(C)** A gene ontology (GO) analysis of Biological Process term enrichment for genes that become significantly derepressed (p-adj < 0.05 and > 1.5-fold) in *Bap1*^*fl/fl*^ cells following OHT treatment. **(D)** Snapshots of Polycomb target genes that become significantly derepressed following BAP1 removal, showing gene expression (cnRNA-seq) and cChIP-seq for H2AK119ub1, H3K27me3, RING1B (PRC1) and SUZ12 (PRC2) in *Bap1*^*fl/fl*^ ESCs (untreated and OHT-treated). **(E)** Heatmaps of cChIP-seq signal for H2AK119ub1, H3K27me3, RING1B (PRC1) and SUZ12 (PRC2) across Polycomb chromatin domains at the promoters of Polycomb target genes in *Bap1*^*fl/fl*^ ESCs (untreated and OHT-treated). Intervals were sorted by RING1B occupancy in untreated *Bap1*^*fl/fl*^ ESCs. **(F)** Metaplots of H2AK119ub1 cChIP-seq signal in *Bap1*^*fl/fl*^ ESCs (untreated and OHT-treated) at the promoters of Polycomb target genes that become significantly derepressed (Up, n = 421) or do not change in expression (No Change, n = 4075) following BAP1 removal. **(G)** As in (F) for H3K27me3 cChIP-seq. **(H)** As in (F) for RING1B cChIP-seq. **(I)** As in (F) for SUZ12 cChIP-seq.

To better understand the interplay between BAP1 and the Polycomb system, we investigated the effect that BAP1 removal has on Polycomb chromatin domains by examining the binding of PRC1 (RING1B), PRC2 (SUZ12), and levels of their respective histone modifications (H2AK119ub1 and H3K27me3) by cChIP-seq. This showed that in the absence of BAP1, H2AK119ub1 increased across Polycomb chromatin domains at target gene promoters (Fig. 5E), although the magnitude of this effect was slightly smaller than at other regions of the genome, in agreement with ChromHMM analysis (Fig. 1E). Furthermore, H3K27me3 was also modestly elevated (Fig. 5E), consistent with an essential role for H2AK119ub1 in shaping H3K27me3 at Polycomb target gene promoters (Blackledge et al. 2014; Cooper et al. 2014; Kalb et al. 2014; Illingworth et al. 2015; Blackledge et al. 2020; Tamburri et al. 2020). In contrast, at the subset of Polycomb target genes that become derepressed in the absence of BAP1, the levels of H2AK119ub1 and H3K27me3 at their promoters remained largely unchanged (Fig. 5D,F,G; Supplemental Fig. S5A,D), suggesting that re-activation of these genes following BAP1 removal is not due to reductions in these histone modifications. We then examined PRC1 and PRC2 occupancy at Polycomb target gene promoters and found that it was modestly reduced, despite the observed increases in H2AK119ub1 and H3K27me3 (Fig. 5E), and this was not due to reductions in RING1B and SUZ12 protein levels (Supplemental Fig. S5B). However, strikingly, when we focused on the promoters of Polycomb target genes that were derepressed in the absence of BAP1, we found that they were on average occupied by much higher levels of PRC1 and PRC2 in untreated cells and showed much more dramatic reductions in their occupancy after BAP1 removal (Fig. 5D,H,I; Supplemental Fig. S5A,C,D). Based on these observations, we conclude that this subset of Polycomb target genes are particularly reliant on high-level occupancy of PRC1 and PRC2 for their silencing, and that the major decrease in Polycomb complex binding at their promoters caused by removal of BAP1 leads to their derepression. Given that both Polycomb repressive complexes can directly bind to H2AK119ub1 (Arrigoni et al. 2006; Zhao et al. 2020; Kalb et al. 2014; Cooper et al. 2016; Kasinath et al. 2020), we envisage that the reductions in PRC1 and PRC2 occupancy at this subset of genes are likely caused by elevated H2AK119ub1 elsewhere in the genome competing for their binding. Together, these findings provide a molecular rationale for the counterintuitive observation that disruption of BAP1/Calypso gives rise to PcG phenotypes in genetic assays, despite their role in counteracting H2AK119ub1. Furthermore, it reveals that limiting pervasive H2AK119ub1 throughout the genome is important for focusing Polycomb repressive complexes at target gene promoters, while enabling transcription elsewhere in the genome.

## Discussion

Chromatin-modifying enzymes can function at defined gene regulatory elements to support cell type-specific gene expression patterns (Atlasi and Stunnenberg 2017; Yadav et al. 2018). Their recruitment to these sites often relies on DNA- and chromatin-binding activities (Smith and Shilatifard 2010), and these mechanisms underpin how PRC1 creates high-level enrichment of H2AK119ub1 at Polycomb target gene promoters to enable repression (Endoh et al. 2012; Blackledge et al. 2015; Fursova et al. 2019; Scelfo et al. 2019; Blackledge et al. 2020; Tamburri et al. 2020; Cohen et al. 2020). In addition to this punctate pool of H2AK119ub1, we and others have recently discovered that PRC1 also places low levels of H2AK119ub1 broadly throughout the genome (Lee et al. 2015; Kahn et al. 2016; Fursova et al. 2019). However, whether pervasive H2AK119ub1 contributes to gene regulation or other chromosomal processes has remained unclear. Here, we discover that BAP1 plays a central role in counteracting pervasive H2AK119ub1, and in its absence, accumulation of H2AK119ub1 throughout the genome leads to widespread reductions in transcription initiation from gene regulatory elements. This reveals an important and previously underappreciated mechanism for chromatin-based gene regulation, whereby a pervasive histone modification can broadly control the function of gene regulatory elements without the need for elaborate site-specific targeting mechanisms.

We envisage that this generic mode of gene regulation could be particularly relevant during cellular differentiation and development when the transcriptional activity of the genome or large genomic regions needs to be coordinately modulated to support acquisition and maintenance of cell type-specific transcriptional states. In fact, support for this concept has recently emerged from studies of X-inactivation where H2AK119ub1 was shown to accumulate across the entire silenced X chromosome to drive transcriptional repression and enable dosage compensation (Fursova et al. 2019; Nesterova et al. 2019; Bousard et al. 2019; Żylicz et al. 2019). Furthermore, our discoveries also indicate that there exists an important balance between the enzymes that place and remove pervasive H2AK119ub1, with the levels of this histone modification regulating the capacity of the genome to be transcribed. Given that the composition and expression of PRC1 and BAP1 complexes changes extensively during development (Fisher et al. 2006; Morey et al. 2012; O’Loghlen et al. 2012; Morey et al. 2015; Kloet et al. 2016), in future work, it will be interesting to investigate how the balance between these two opposing activities is regulated at different developmental stages and how cell type-specific H2AK119ub1 levels influence the transcription potential of the genome. Given that BAP1 and other PR-DUB subunits are frequently mutated in a variety of cancers with diverse origins (Wiesner et al. 2011; Dey et al. 2012; Carbone et al. 2013; Murali et al. 2013; Katoh 2013; Masoomian et al. 2018; Zhang et al. 2020), our findings also suggest that maintaining the cell type-specific balance between the activities that control H2AK119ub1 levels could play an important role in protecting cells from transformation.

BAP1 has previously been proposed to regulate gene expression through diverse mechanisms, some of which are thought to function independently of H2AK119ub1 (Yu et al. 2010; Dey et al. 2012; Li et al. 2017b; Wang et al. 2018; Kuznetsov et al. 2019; Campagne et al. 2019). We now discover that BAP1 plays a surprisingly widespread role in supporting gene expression and show that this relies on BAP1 counteracting H2AK119ub1, as catalytic inactivation of PRC1 reverts the effects of BAP1 removal on gene expression. This raises the important question of how pervasive H2AK119ub1 can function at such a broad scale to constrain gene expression. Many histone modifications are thought to regulate gene expression through reader proteins which bind to modified nucleosomes and directly affect transcription (Musselman et al. 2012; Patel and Wang 2013). This is particularly relevant for histone modifications that are of low abundance yet highly enriched at gene regulatory elements. However, we estimate that approximately 5.9×10^6^ H2AK119ub1 molecules decorate the genome of ES cells (Huseyin and Klose 2020), and this number nearly doubles in the absence of BAP1. If a reader protein was required for the widespread repressive effects of H2AK119ub1, we envisage that it would also need to be immensely abundant, a requirement that none of the proposed H2AK119ub1-binding proteins fulfil (Arrigoni et al. 2006; Richly et al. 2010; Kalb et al. 2014; Qin et al. 2015; Cooper et al. 2016; Zhang et al. 2017b; Beck et al. 2011; Schwanhäusser et al. 2011; Wiśniewski et al. 2014). Alternatively, addition of a bulky ubiquitin moiety to histone H2A could simply restrict access of the transcriptional machinery to gene regulatory elements. However, in agreement with previous work (Hodges et al. 2018; King et al. 2018), we find that accumulation of H2AK119ub1 does not limit chromatin accessibility, and if anything, the genome becomes slightly more accessible when H2AK119ub1 levels are increased. Based on these observations, we favor the possibility that H2AK119ub1 controls gene expression by counteracting the process of transcription more directly. In agreement with this, it has previously been suggested that PRC1 and H2AK119ub1 can inhibit various aspects of transcription, including initiation, pause release and elongation (Dellino et al. 2004; Stock et al. 2007; Zhou et al. 2008; Nakagawa et al. 2008; Lehmann et al. 2012; Aihara et al. 2016). By examining how elevated H2AK119ub1 affects Pol II, we now discover that it preferentially compromises Ser5-phosphorylation at gene regulatory elements, suggesting that H2AK119ub1 acts primarily through counteracting transcription initiation. Consistent with this, we have recently shown that rapid depletion of H2AK119ub1 leads to increased Polycomb target gene expression which results from increased transcription initiation (Dobrinić et al. 2020). These findings are also in agreement with observations from *in vitro* studies where installation of H2AK119ub1 into chromatin templates was sufficient to impede transcription initiation (Nakagawa et al. 2008). Therefore, pervasive H2AK119ub1 appears to regulate the function of gene regulatory elements by limiting transcription initiation.

Our molecular understanding of how BAP1 and its H2AK119ub1 deubiquitylase activity contribute to gene regulation has been complicated by the initial characterisation of BAP1 as a PcG gene in genetic experiments in *Drosophila* (de Ayala Alonso et al. 2007; Scheuermann et al. 2010). Given that PcG genes are known to maintain Polycomb target gene repression during development, it was puzzling why BAP1, which removes H2AK119ub1, would be required for this process. Initially, it was proposed that BAP1 functioned at Polycomb target gene promoters to appropriately balance H2AK119ub1 levels by enabling its dynamic turnover, and that this would somehow facilitate repression of these genes (Scheuermann et al. 2010; Schuettengruber and Cavalli 2010). While alternative and less direct mechanisms have also been previously considered (Schuettengruber and Cavalli 2010; Scheuermann et al. 2012; Gutiérrez et al. 2012), here we demonstrate that BAP1 indirectly supports repression of a subset of Polycomb target genes by limiting pervasive H2AK119ub1 elsewhere in the genome to promote high-level occupancy of Polycomb complexes at target gene promoters. Interestingly, mutations in the BAP1 and ASXL components of the PR-DUB complex can also lead to phenotypes that are reminiscent of mutations in Trithorax group (TrxG) genes, which are known to oppose PcG gene activity and facilitate gene expression (Sinclair et al. 1992; Milne et al. 1999; Gildea et al. 2000; Baskind et al. 2009; Scheuermann et al. 2010; Fisher et al. 2010). In line with these observations, we demonstrate at the molecular level that by counteracting pervasive H2AK119ub1, BAP1 directly promotes gene expression, akin to a TrxG gene (Fig. 6A), while also indirectly supporting repression of a subset of Polycomb target genes, akin to a PcG gene (Fig. 6B). Together, these new discoveries provide a mechanistic explanation for the dual role that BAP1 has in gene regulation based on genetic assays, and reveals that the balance between activities that place and remove pervasive H2AK119ub1 is essential for supporting the expression of some genes, while maintaining the repression of others.

**Figure 6.**
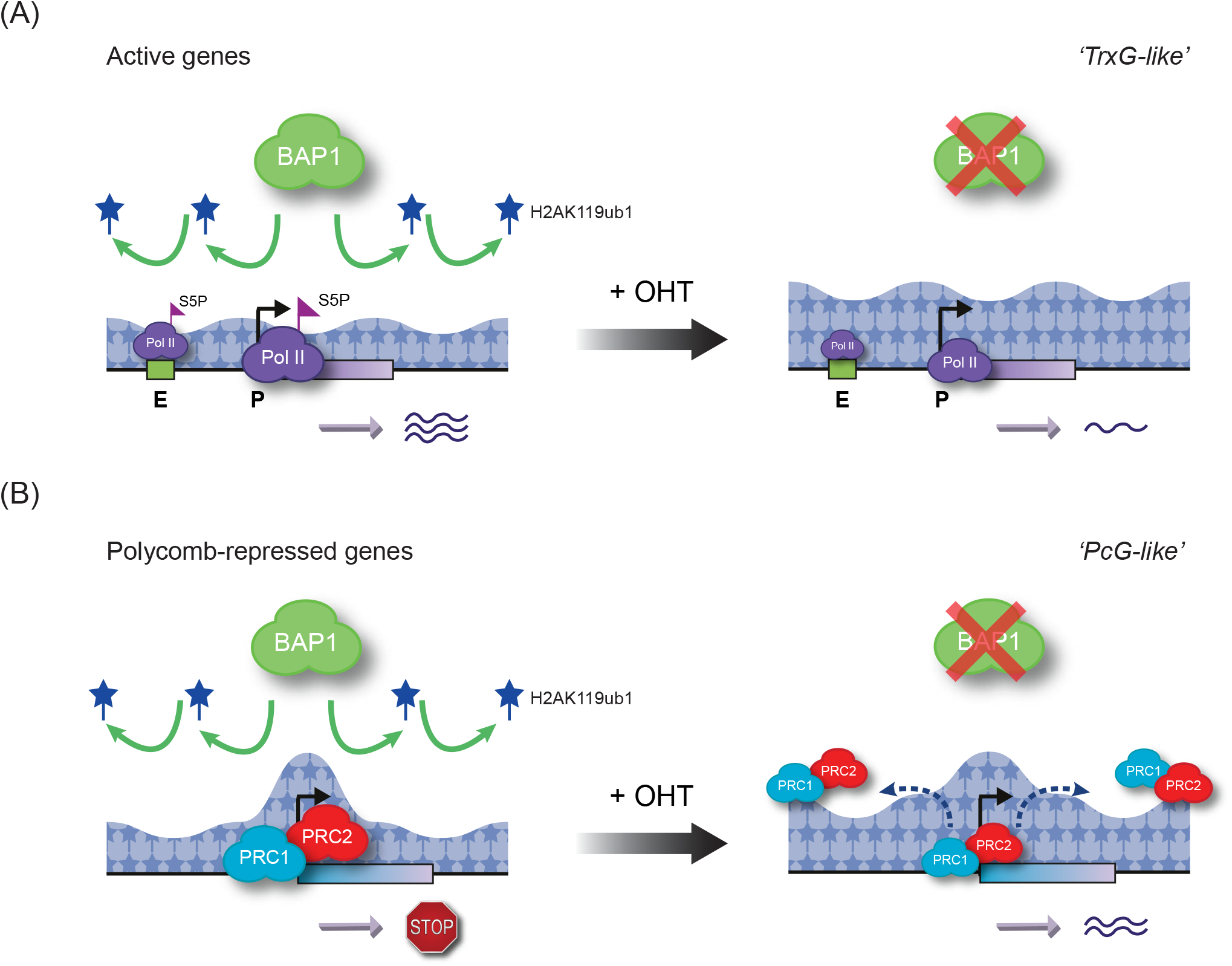
A model illustrating how BAP1 can regulate gene expression by constraining pervasive H2AK119ub1. **(A)** BAP1 facilitates gene expression by constraining the pervasive sea of H2AK119ub1 that covers the genome. Inducible removal of BAP1 (+ OHT) results in a broad accumulation of H2AK119ub1 throughout the genome. Elevated H2AK119ub1 indiscriminately counteracts transcription initiation-associated Ser5-phosphorylation (S5P) of Pol II at gene regulatory elements (P=promoters/E=enhancers), which leads to widespread reductions in transcription and gene expression. This explains why disruption of BAP1 and other PR-DUB subunits can lead to Trithorax group (TrxG)-like phenotypes. **(B)** BAP1 also indirectly supports repression of a subset of Polycomb target genes by counteracting pervasive H2AK119ub1 and focusing high levels of Polycomb complexes at target gene promoters. In the absence of BAP1, PRC1/PRC2 occupancy at Polycomb target sites is reduced, presumably due to the increased binding of these complexes to elevated H2AK119ub1 elsewhere in the genome. This leads to derepression of a subset of Polycomb target genes that appear to rely on high-level Polycomb complex occupancy for their silencing, explaining why BAP1 was originally characterised as a Polycomb group (PcG) gene.

## Acknowledgments

Work in the Klose lab is supported by the Wellcome Trust (209400/Z/17/Z), the European Research Council (681440), and the Lister Institute of Preventive Medicine. We would like to thank Amanda Williams at the Department of Zoology, Oxford, for sequencing support on the NextSeq 500. We thank Dr Emilia Dimitrova for critical reading of the manuscript.

## Author contributions

Conceptualization, N.A.F., N.P.B. and R.J.K.; Methodology, N.A.F., A.H.T., N.P.B.; Investigation, N.A.F., A.H.T., E.L.F., N.P.B., and P.D.; Formal Analysis, Software and Data Curation, N.A.F; Resources, N.A.F., A.L., N.P.B., and M.K.H.; Writing – Original Draft, N.A.F., and R.J.K.; Writing – Review and Editing, N.A.F. and R.J.K., with the input from all the other authors; Funding Acquisition, R.J.K.; Supervision, R.J.K.

## Materials and Methods

### Cell culture conditions and treatments

E14TG2a mouse embryonic stem cells (ESCs) were grown on gelatin-coated plates at 37°C and 5% CO2 in Dulbecco’s Modified Eagle Medium (DMEM) supplemented with 15% fetal bovine serum (Labtech), 2 mM L-glutamine (Life Technologies), 1x penicillin/streptomycin (Life Technologies), 1x non-essential amino acids (Life Technologies), 0.5 mM beta-mercaptoethanol (Life Technologies), and 10 ng/ml leukemia inhibitory factor (in-house). To induce conditional removal of BAP1, on its own or in combination with PRC1 catalytic activity, *Bap1*^*fl/fl*^ and *PRC1*^*CPM*^;*Bap1*^*fl/fl*^ cells were treated with 800 nM 4-hydroxytamoxifen (OHT) for 96 hr. To induce conditional removal of PRC1 catalytic activity on its own, *PRC1*^*CPM*^ cells were treated with 800 nM OHT for 72 hr. Cells were regularly tested for the presence of mycoplasma.

Human HEK293T cells used for spike-in calibration of cChIP-seq were grown at 37°C and 5% CO2 in Dulbecco’s Modified Eagle Medium (DMEM) supplemented with 10% fetal bovine serum (Labtech), 2 mM L-glutamine (Life Technologies), 1x penicillin-streptomycin (Life Technologies), and 0.5 mM beta-mercaptoethanol (Life Technologies). *Drosophila* S2 (SG4) cells used for spike-in calibration of Native cChIP-seq, cnRNA-seq and cATAC-seq were grown adhesively at 25°C in Schneider’s Drosophila Medium (Life Technologies), supplemented with 1x penicillin-streptomycin (Life Technologies) and 10% heat-inactivated fetal bovine serum (Labtech).

### Cell line generation

*Bap1*^*fl/fl*^ cells were derived from E14TG2a ESCs by a two-step process. First, parallel loxP sites flanking exon 4 of the *Bap1* gene were inserted using a targeting construct with homology arms of approximately 1 kb and three different Cas9 guides specific for the *Bap1* locus (sgRNA target sequences: TCAAATGGATCGAAGAGCGC, CAAGGTAGGGACACAATAAA, TAAAACACCACCAACTACAG). Second, CreERT2 was inserted into the *Rosa26* locus using a Rosa26-specific Cas9 guide (sgRNA target sequence: CGCCCATCTTCTAGAAAGAC). The same targeting construct and Cas9 guides were used to generate *PRC1*^*CPM*^;*Bap1*^*fl/fl*^ cells from a *PRC1*^*CPM*^ parental cell line. Loss of BAP1 in response to OHT treatment in *Bap1*^*fl/fl*^ and *PRC1*^*CPM*^;*Bap1*^*fl/fl*^ ESCs was confirmed using RT-qPCR and western blot analysis.

*PRC1*^*CPM*^ cells were generated and characterised in a previous study (Blackledge et al. 2020). Briefly, a targeting construct for this cell line comprised exon 3 of *Ring1b* in forward orientation (flanked by 100 bp of *Ring1b* intron 2/intron 3), followed by a mutant copy of exon 3 (encoding I53A and D56K mutations) in reverse orientation (flanked by splice donor and acceptor sites from mouse *IgE* gene). Both the wild-type and mutant versions of exon 3 were codon-optimized at wobble positions to minimize sequence similarity. The wild-type/mutant exon 3 pair was flanked by doubly inverted LoxP/Lox2272 sites and approximately 1 kb homology arms. The targeting construct was transfected into E14TG2a ESCs in combination with three different Cas9 guides specific for the *Ring1b* gene. Correctly targeted homozygous clones were identified by PCR screening, followed by RT-PCR and sequencing to check for splicing defects. Using a similar approach, the I50A/D53K mutation was constitutively knocked-in into both copies of the endogenous *Ring1a* gene. Finally, CreERT2 was inserted into the *Rosa26* locus using a Rosa26-specific Cas9 guide.

### Genome engineering by CRISPR/Homology-Directed Repair (HDR)

The pSptCas9(BB)-2A-Puro(PX459)-V2.0 vector was obtained from Addgene (#62988) and sgRNAs were designed using the CRISPOR online tool (http://crispor.tefor.net/crispor.py). Targeting constructs with appropriate homology arms were generated by Gibson assembly using the Gibson Assembly Master Mix kit (NEB). Targeting constructs were designed such that Cas9 recognition sites were disrupted by the presence of the LoxP sites. ESCs (one well of a 6-well plate) were transfected with 0.5 μg of each Cas9 guide and 2 μg of targeting construct using Lipofectamine 3000 (ThermoFisher) according to manufacturer’s guidelines. The day after transfection, cells were passaged at a range of densities and subjected to selection with 1 μg/ml puromycin for 48 hr to eliminate non-transfected cells. Approximately a week later, individual clones were isolated, expanded and PCR-screened for the desired genomic modifications.

### Preparation of nuclear and histone extracts for immunoblotting analysis

For nuclear extraction, ESCs were washed with 1x PBS and resuspended in 10 volumes of Buffer A (10 mM Hepes pH 7.9, 1.5 mM MgCl_2_, 10 mM KCl, 0.5 mM DTT, 0.5 mM PMSF and 1x protease inhibitor cocktail (PIC) (Roche)). After 10 min incubation on ice, cells were recovered by centrifugation at 1500 g for 5 min and resuspended in 3 volumes of Buffer A supplemented with 0.1% NP-40. The released nuclei were pelleted by centrifugation at 1500 g for 5 min, followed by resuspension in 1 volume of Buffer C (5 mM Hepes pH 7.9, 26% glycerol, 1.5 mM MgCl_2_, 0.2 mM EDTA, 1xPIC (Roche) and 0.5 mM DTT) supplemented with 400 mM NaCl. The extraction was allowed to proceed on ice for 1 hr with occasional agitation, then the nuclei were pelleted by centrifugation at 16,000 g for 20 min at 4°C. The supernatant was taken as the nuclear extract. The Bradford protein assay (BioRad) was used to compare protein concentrations across samples.

For histone extraction, ESCs were washed in RSB (10 mM Tris HCl pH 8, 10 mM NaCl, 3 mM MgCl_2_) supplemented with 20 mM N-ethylmaleimide (NEM), incubated on ice for 10 min in RSB with 0.5% NP-40 and 20 mM NEM, pelleted by centrifugation at 800 g for 5 min, and incubated in 2.5 mM MgCl_2_, 0.4 M HCl and 20 mM NEM on ice for 30 min. After that, cells were pelleted by centrifugation at 16,000 g at 4°C for 20 min, the supernatant recovered and precipitated on ice with 25% TCA for 30 min, followed by centrifugation at 16,000 g for 15 min at 4°C to recover histones. Following two acetone washes, the histones were resuspended in 1x SDS loading buffer (2% SDS, 100 mM Tris pH 6.8, 100 mM DTT, 10% glycerol and 0.1% bromophenol blue) and boiled at 95°C for 5 min. Finally, any insoluble precipitate was pelleted by centrifugation at 16,000 g for 10 min, and the soluble fraction retained as the histone extract. Histone concentrations across samples were compared using SDS-PAGE followed by Coomassie Blue staining. Western blot analysis of nuclear and histone extracts was performed using LI-COR IRDye secondary antibodies, and imaging was done using the LI-COR Odyssey Fc system. A list of antibodies used in this study for western blot and cChIP-seq analysis can be found in Supplemental Table S1.

**Supplemental Table S1.** A list of antibodies used in this study for western blot and cChIP-seq analysis.

| Antibody | Source | Identifier | Application |
| --- | --- | --- | --- |
| Rabbit monoclonal anti-H2AK119ub1 | Cell Signalling Technology | Cat#:8240 | Western and ChIP |
| Rabbit monoclonal anti-H2BK120ub1 | Cell Signalling Technology | Cat#:5546 | Western |
| Rabbit polyclonal anti-H3K27me3 | In house (Rose et al., 2016) | N/A | ChIP |
| Rabbit polyclonal anti-H3K4me3 | In house (Farcas et al., 2012) | N/A | ChIP |
| Rabbit monoclonal anti-H3K27ac | Cell Signalling Technology | Cat#:8173 | ChIP |
| Rabbit monoclonal anti-H3K4me1 | Cell Signalling Technology | Cat#:5326 | ChIP |
| Mouse monoclonal anti-H3 | Cell Signalling Technology | Cat#:3638 | Western |
| Rabbit monoclonal anti-RING1B | Cell Signalling Technology | Cat#:5694 | ChIP |
| Mouse monoclonal anti-RING1B | In house (Atsuta et al., 2001) | N/A | Western |
| Rabbit monoclonal anti-SUZ12 | Cell Signalling Technology | Cat#:3737 | Western and ChIP |
| Rabbit monoclonal anti-BAP1 | Cell Signalling Technology | Cat#:13271 | Western |
| Rabbit monoclonal anti-ASXL1 | Cell Signalling Technology | Cat#:52519 | Western |
| Rabbit polyclonal anti-FOXK1 | Cell Signalling Technology | Cat#:12025 | Western |
| Rabbit monoclonal anti-BRG1 | Abcam | ab110641 | Western |
| Rabbit monoclonal anti-Rpb1-NTD | Cell Signalling Technology | Cat#:14958 | ChIP |
| Rabbit monoclonal anti-Rpb1-S5P | Cell Signalling Technology | Cat#:13523 | ChIP |
| Rabbit monoclonal anti-Rpb1-S2P | Cell Signalling Technology | Cat#:13499 | ChIP |

### Calibrated ChIP-sequencing (cChIP-seq)

For RING1B and SUZ12, cChIP-seq was performed as described previously (Fursova et al. 2019; Blackledge et al. 2020). Briefly, 5×10^7^ mouse ESCs (untreated and OHT-treated) were crosslinked in 10 ml 1x PBS with 2 mM DSG (Thermo Scientific) for 45 minutes at 25°C, and then with 1% formaldehyde (methanol-free, Thermo Scientific) for a further 15 minutes. Crosslinking was stopped by quenching with 125 mM glycine. Crosslinked ESCs were mixed with 2×10^6^ human HEK293T cells, which have been similarly double-crosslinked, and incubated in lysis buffer (50 mM HEPES pH 7.9, 140 mM NaCl, 1 mM EDTA, 10% glycerol, 0.5% NP40, 0.25% Triton-X100, 1x PIC (Roche)) for 10 minutes at 4°C. Released nuclei were washed (10 mM Tris-HCl pH 8, 200 mM NaCl, 1 mM EDTA, 0.5 mM EGTA, 1x PIC (Roche)) for 5 minutes at 4°C. Chromatin was then resuspended in 1 ml of sonication buffer (10 mM Tris-HCl pH 8, 100 mM NaCl, 1 mM EDTA, 0.5 mM EGTA, 0.1% Na deoxycholate, 0.5% N-lauroylsarcosine, 1x PIC (Roche)) and sonicated for 30 minutes using the BioRuptor Pico (Diagenode), shearing genomic DNA to an average size of 0.5 kb. Following sonication, TritonX-100 was added to a final concentration of 1%, followed by centrifugation at 20,000 g for 10 min at 4°C to collect the clear supernatant fraction.

For Pol II and its phosphorylated forms, cChIP-seq was done as described previously (Turberfield et al. 2019). Briefly, 5×10^7^ ESCs (untreated and OHT-treated) were crosslinked in 10 ml 1x PBS with 1% formaldehyde (methanol-free, Thermo Scientific) for 10 min at 25°C and then quenched by addition of 125 mM glycine. Cross-linked ESCs were mixed with 2×106 human HEK293T cells, which have been similarly single-crosslinked, and then incubated in FA-lysis buffer (50 mM HEPES pH 7.9, 150 mM NaCl, 2 mM EDTA, 0.5 mM EGTA, 0.5% NP40, 0.1% sodium deoxycholate, 0.1% SDS, 10 mM NaF, 1 mM AEBSF, 1×PIC) for 10 min at 4°C. Chromatin was sonicated for 30 min using the BioRuptor Pico (Diagenode), followed by centrifugation at 20,000 g for 10 min at 4°C to collect the clear supernatant fraction.

For RING1B and SUZ12 ChIP, sonicated chromatin was diluted 10-fold with ChIP dilution buffer (1% Triton-X100, 1 mM EDTA, 20 mM Tris-HCl pH 8, 150 mM NaCl, 1xPIC). For Pol II ChIP, 300 ug of chromatin per one IP was diluted in FA-lysis buffer up to a final volume of 1 ml. Diluted chromatin was pre-cleared for 1 hr using Protein A agarose beads (Repligen) that were pre-blocked with 1 mg/ ml BSA and 1 mg/ml yeast tRNA. For each ChIP reaction, 1 ml of diluted and pre-cleared chromatin was incubated overnight with the appropriate antibody, anti-RING1B (CST, D22F2, 3 μl), anti-SUZ12 (CST, D39F6, 3 μl), anti-Rpb1-NTD (CST, D8L4Y, 15 μl) as a measure of total Pol II levels, anti-Rpb1-CTD-Ser5P (CST, D9N5I, 12.5 μl) and anti-Rpb1-CTD-Ser2P (CST, E1Z3G, 12.5 μl) as a measure of Pol II phosphorylation levels. To capture antibody-bound chromatin, ChIP reactions were incubated with pre-blocked protein A agarose beads (Repligen) for 2 hr (RING1B and SUZ12) or 3 hr (Pol II) at 4°C. For RING1B and SUZ12, ChIP washes were performed as described previously (Farcas et al. 2012). For Pol II, washes were performed with FA-Lysis buffer, FA-Lysis buffer containing 500 mM NaCl, DOC buffer (250 mM LiCl, 0.5% NP-40, 0.5% sodium deoxycholate, 2 mM EDTA, 10 mM Tris-HCl pH 8), followed by two washes with TE buffer (pH 8). ChIP DNA was eluted in elution buffer (1% SDS, 0.1 M NaHCO3) and cross-linking was reversed overnight at 65°C with 200 mM NaCl and 2 μl RNase A (Sigma). Matched input samples (10% of original ChIP reaction) were treated identically. The following day, ChIP samples and inputs were incubated with Proteinase K (Sigma) for at least 1.5 hr at 56°C and purified using ChIP DNA Clean and Concentrator Kit (Zymo Research).

cChIP-seq libraries for both ChIP and Input samples were prepared using NEBNext Ultra II DNA Library Prep Kit for Illumina, following manufacturer’s guidelines. Samples were indexed using NEBNext Multiplex Oligos. The average size and concentration of all libraries were analysed using the 2100 Bioanalyzer High Sensitivity DNA Kit (Agilent) followed by qPCR quantification using SensiMix SYBR (Bioline, UK) and KAPA Illumina DNA standards (Roche). Libraries were sequenced as 40 bp paired-end reads in biological triplicate or quadruplicate on Illumina NextSeq 500 platform.

### Native cChIP-sequencing

Native cChIP-seq for H2AK119ub1, H3K27me3, H3K27ac, H3K4me3, and H3K4me1 was performed as described previously (Fursova et al. 2019). Briefly, 5×10^7^ mouse ESCs (untreated and OHT-treated) were mixed with 2×10^7^ *Drosophila* SG4 cells in 1x PBS. Mixed cells were pelleted and nuclei were released by resuspending in ice-cold lysis buffer (10 mM Tris-HCl pH 8, 10 mM NaCl, 3 mM MgCl_2_, 0.1% NP40, 5 mM sodium butyrate, and 5 mM N-ethylmaleimide). Nuclei were then washed and resuspended in 1 ml of MNase digestion buffer (10 mM Tris-HCl pH 8.0, 10 mM NaCl, 3 mM MgCl_2_, 0.1% NP40, 0.25 M sucrose, 3 mM CaCl_2_, 10 mM sodium butyrate, 10 mM N-ethylmaleimide, and 1x PIC (Roche)). Each sample was incubated with 200 units of MNase (Fermentas) at 37°C for 5 min, followed by the addition of 4 mM EDTA to halt MNase digestion. Following centrifugation at 1500 g for 5 min at 4°C, the supernatant (S1) was retained. The remaining pellet was incubated with 300 μl of nucleosome release buffer (10 mM Tris-HCl pH 7.5, 10 mM NaCl, 0.2 mM EDTA, 10 mM sodium butyrate, 10 mM N-ethylmaleimide, and 1x PIC (Roche)) at 4°C for 1 hr, passed five times through a 27G needle using a 1 ml syringe, and spun at 1500 g for 5 min at 4°C. The second supernatant (S2) was collected and combined with the corresponding S1 sample from above. Digestion to mostly mono-nucleosomes was confirmed by agarose gel electrophoresis of purified S1/S2 DNA.

For ChIP, S1/S2 nucleosomes were diluted 10-fold in Native ChIP incubation buffer (70 mM NaCl, 10 mM Tris pH 7.5, 2 mM MgCl_2_, 2 mM EDTA, 0.1% TritonX-100, 10 mM sodium butyrate (for H3K27ac and H3K4me3 ChIPs), 10 mM N-ethylmaleimide, and 1xPIC (Roche)). Each ChIP reaction, 1 ml of diluted nucleosomes was incubated overnight at 4°C with the appropriate antibody, anti-H2AK119ub1 (CST, D27C4, 5 μl), anti-H3K27me3 (in-house, 5 μl), anti-H3K27ac (CST, D5E4, 3 μl), anti-H3K4me3 (in-house, 4 μl) or anti-H3K4me1 (CST, D1A9, 5 μl). Antibody-bound nucleosomes were captured for 1 hr at 4°C using protein A agarose (Repligen) beads, pre-blocked in Native ChIP incubation buffer supplemented with 1 mg/ml BSA and 1 mg/ml yeast tRNA, and collected by centrifugation. Immunoprecipitated material was washed four times with Native ChIP wash buffer (20 mM Tris pH 7.5, 2 mM EDTA, 125 mM NaCl, 0.1% Triton-X100) and once with TE buffer (pH 8). ChIP DNA was eluted using 100 μl of elution buffer (1% SDS, 0.1 M NaHCO3) for 30 min at room temperature, and then purified using ChIP DNA Clean and Concentrator Kit (Zymo Research). For each ChIP sample, DNA from a matched input control (corresponding to 10% of original ChIP reaction) was purified in the same way. Native cChIP-seq library preparation and sequencing was performed as described above for cChIP-seq.

### Calibrated nuclear RNA-sequencing (cnRNA-seq)

For cnRNA-seq, 1×10^7^ ESCs (untreated and OHT-treated) were mixed with 4×10^6^ *Drosophila* SG4 cells in 1x PBS. Nuclei were isolated in 1 ml HS Lysis buffer (50 mM KCl, 10 mM MgSO_4_.7H_2_0, 5 mM HEPES, 0.05% NP40 (IGEPAL CA630), 1 mM PMSF, 3 mM DTT, 1xPIC (Roche)) for 1 min at room temperature, and then recovered by centrifugation at 1000 g for 5 min at 4°C, followed by a total of three washes with ice-cold RSB buffer (10 mM NaCl, 10 mM Tris pH 8, 3 mM MgCl_2_). Nuclei integrity was assessed using 0.4% Trypan Blue staining (ThermoScientific). Pelleted nuclei were resuspended in 1 ml of TRIzol reagent (ThermoScientific), and RNA was extracted according to the manufacturer’s protocol, followed by treatment with the TURBO DNA-free Kit (ThermoScientific) to remove any contaminating DNA. Quality of RNA was assessed using the 2100 Bioanalyzer RNA 6000 Pico kit (Agilent). RNA samples were depleted of rRNA with the NEBNext rRNA Depletion kit (NEB) prior to preparing cnRNA-seq libraries using the NEBNext Ultra (for *Bap1*^*fl/fl*^ and *PRC1*^*CPM*^ ESCs) or Ultra II (for *PRC1*^*CPM*^;*Bap1*^*fl/fl*^ ESCs) Directional RNA Library Prep kit (NEB). To quantitate the consistency of spike-in cell mixing for each individual sample, a small aliquot of nuclei was saved to isolate genomic DNA using phenol-chloroform extraction. This was followed by sonication of DNA for 15 min using the BioRuptor Pico (Diagenode), shearing genomic DNA to an average size of less than 1 kb. Libraries from sonicated genomic DNA were constructed as described above for cChIP-seq. Both cnRNA-seq and gDNA-seq libraries were sequenced as 80 bp paired-end reads on the Illumina NextSeq 500 platform in biological triplicates.

### Calibrated ATAC-seq (cATAC-seq)

To assay chromatin accessibility, calibrated ATAC-seq was performed as described previously (Turberfield et al. 2019). First, 1×10^7^ ESCs (untreated and OHT-treated) were mixed with 4×10^6^ *Drosophila* SG4 cells in 1x PBS and then lysed in 1 ml HS Lysis buffer (50 mM KCl, 10 mM MgSO_4_.7H_2_0, 5 mM HEPES, 0.05% NP40 (IGEPAL CA630), 1 mM PMSF, 3 mM DTT, 1xPIC (Roche)) for 1 min at room temperature. Nuclei were recovered by centrifugation at 1000 g for 5 min at 4°C and washed three times in ice-cold RSB buffer (10 mM NaCl, 10 mM Tris pH 7.4, 3 mM MgCl_2_). The concentration and integrity of nuclei were assessed using 0.4% Trypan Blue staining (ThermoScientific). Next, 5×10^5^ nuclei were resuspended in Tn5 reaction buffer (10 mM TAPS, 5 mM MgCl2, 10% dimethylformamide) and incubated with Tn5 transposase (25 μM, generated in house as previously described (King and Klose 2017)) for 30 min at 37°C. Tagmented DNA was purified using MinElute columns (QIAGEN) and eluted in 10 μl elution buffer. To control for the Tn5 transposase sequence bias and to determine the exact spike-in ratio for each individual sample, 50 ng of genomic DNA, isolated from the same nuclei preparation by phenol-chloroform extraction, was tagmented with Tn5 transposase (25 μM) for 30 min at 55°C and purified using MinElute columns (QIAGEN).

Libraries for cATAC-seq and gDNA-seq were prepared by PCR amplification using the NEBNext High-Fidelity 2X PCR Master Mix and custom-made Illumina barcodes (Buenrostro et al. 2015). Libraries were purified by two rounds of Agencourt AMPure XP bead cleanup (1.5× bead:sample ratio). Library concentration and fragment size distribution were determined as described above for cChIP-seq. Libraries were sequenced using the Illumina NextSeq 500 platform in biological quadruplicate using 40 bp paired-end reads.

### Data processing and normalisation for massive parallel sequencing

Following massive parallel sequencing, reads were mapped and processed as described previously (Fursova et al. 2019; Turberfield et al. 2019). Briefly, for cChIP-seq, cATAC-seq and gDNA-seq, Bowtie 2 (Langmead and Salzberg 2012) was used to align paired-end reads to the concatenated mouse and spike-in genome sequences (mm10+dm6 for native cChIP-seq, cATAC-seq and gDNA-seq; mm10+hg19 for cross-linked cChIP-seq) with the “--no-mixed” and “--no-discordant” options. Non-uniquely mapping reads were discarded, and PCR duplicates were removed with Sambamba (Tarasov et al., 2015). For cATAC-seq, reads mapping to the mitochondrial chromosome and other genomic regions with artificially high counts or low mappability, derived from the ENCODE blacklist (Amemiya et al. 2019), were also discarded.

For cnRNA-seq, first, Bowtie 2 was used with “--very-fast”, “--no-mixed” and “--no-discordant” options to identify and discard reads mapping to the concatenated mm10+dm6 rDNA genomic sequences (GenBank: BK000964.3 and M21017.1). Next, all unmapped reads were aligned against the concatenated mm10+dm6 genome using STAR (Dobin et al. 2013). To improve mapping of intronic sequences, reads that failed to be mapped by STAR were further aligned with Bowtie 2 (with “--sensitive-local”, “--no-mixed” and “--no-discordant” options). Uniquely aligned reads from the last two steps were combined, and PCR duplicates were removed using Sambamba (Tarasov et al. 2015). A list of all genomics datasets produced in this study and the number of uniquely aligned reads in each experiment can be found in Supplemental Table S2.

To visualise the cChIP-seq, cATAC-seq and cnRNA-seq and quantitatively compare genome enrichment profiles, chromatin accessibility and gene expression between the conditions, the data were internally calibrated using dm6 or hg19 spike-in as described previously (Fursova et al. 2019; Turberfield et al. 2019). Briefly, uniquely aligned mm10 reads were randomly subsampled based on the total number of spike-in (dm6 or hg19) reads in each sample. To account for variations in the spike-in cell mixing, we used the ratio of spike-in/mouse total read counts in the corresponding Input/gDNA-seq samples to correct the subsampling factors. After normalisation, read coverages across genomic regions of interest (RING1B peaks for H2AK119ub1, H3K27me3, RING1B and SUZ12 cChIP-seq, TSS ± 2.5 kb for cATAC-seq and H3K27ac, H3K4me3 and H3K4me1 cChIP-seq, or gene bodies for total Pol II, Ser5P- and Ser2P-Pol II cChIP-seq and cnRNA-seq) were compared for individual biological replicates using multiBamSummary and plotCorrelation from deepTools (Ramírez et al. 2014). For each experimental condition, biological replicates showed a good correlation (Pearson correlation coefficient > 0.9, see Supplemental Table S3) and were merged for downstream analysis. Genome coverage tracks were generated using the pileup function from MACS2 (Zhang et al. 2008) for cChIP-seq, bamCoverage from deeptools (Ramírez et al. 2014) for cATAC-seq, and genomeCoverageBed from BEDTools (Quinlan and Hall 2010) for cnRNA-seq, and visualised using the UCSC genome browser (Kent et al. 2002). BigwigCompare from deeptools was used to make differential genome coverage tracks (log2 ratio of two conditions or ratio of Pol II phosphorylated forms to its total levels).

### Genome segmentation by ChromHMM

ChromHMM (Ernst and Kellis 2012) was used to perform unsupervised segmentation of the genome into distinct chromatin states, enriched with different combinations of histone modifications and other chromatin features, as described previously, but without extension of the reads (Ernst and Kellis 2017). Briefly, to build the model, cATAC-seq and cChIP-seq for H2AK119ub1, H3K27me3, H3K27ac, H3K4me3 and H3K4me1 from this study (GEO:GSEXXXXXX), together with published ChIP-seq datasets for CTCF (GEO:GSE153400, (Huang et al. 2020)), OCT4, NANOG, SOX2 (GEO:GSE87822, (King and Klose 2017)), H3K36me3 and H3K9me3 (GEO:GSE120376, (Ramisch et al. 2019)) in wild-type ESCs, were aligned as described above, subsampled to the same number of reads and binarized using the binarizeBam function in a paired-end mode. These data were used to learn ChromHMM models with 10-15 states using the LearnModel function with default parameters. Finally, a 13-state ChromHMM model was selected for the downstream analysis, as this was the minimal number of states required to accurately segregate an active enhancer state based on the high frequency of OCT4/NANOG/SOX2 binding. This resulted in 13 different chromatin states which were interpreted as described in Supplemental Fig. S1B based on the underlying functional genomic annotations and their transcriptional activity. These included CTCF-bound insulators, distal gene regulatory elements (weak to highly active enhancers), actively transcribed promoters and gene body regions, Polycomb-repressed and bivalent chromatin domains, H3K9me3-enriched heterochromatin and intergenic regions, not enriched with any examined chromatin features.

### Peak calling and annotation of genomic regions

To define active gene regulatory elements, we first performed peak calling for H3K27ac cChIPseq in *Bap1*^*fl/fl*^ ESCs (untreated and OHT-treated) for four biological replicates using the dpeak function (-kd 500, -kw 300, -p 1e-30) from DANPOS2 (Chen et al. 2013), discarding peaks that overlapped with a custom set of blacklisted genomic regions. We then intersected ATAC peaks defined previously (King et al. 2018) with our H3K27ac peak set. This resulted in a set of active gene regulatory elements, which were further segregated into active enhancers (non-TSS, n = 12,006) or active promoters (TSS, n = 10,840), based on their overlap with TSS ± 1 kb regions. To obtain the most complete set of TSS positions, we combined TSS annotations from the UCSC refGene (n = 34,852), NCBI RefSeq (n = 106,520) and GENCODE VM24 (n = 67,573) databases. To eliminate the contribution of gene body signal for Pol II cChIP-seq, only intergenic active enhancers (n = 4156), which did not overlap with any of the above gene annotations, were considered for the downstream analysis. To associate genes with putative distal regulatory elements, for each gene promoter we identified the nearest intergenic enhancer located within the 250 kb distance using the closest function from BEDTools. For differential gene expression analysis and quantification of cChIP-seq and cATAC-seq signal at promoters or over the bodies of genes, we used a custom non-redundant set of genes (n = 20,633), derived from mm10 UCSC refGene genes by removing very short genes with poor sequence mappability and highly similar transcripts as described previously (Rose et al. 2016). For the purposes of read quantification, promoters were defined as TSS ± 2.5 kb intervals, with the exception of Pol II cChIP-seq, in which case they were defined as TSS ± 0.5 kb to specifically capture the promoter signal and eliminate the contribution of the gene body signal. A set of intergenic intervals was obtained using the complement function from BEDTools as regions of the genome that do not overlap with any of the genes from a complete UCSC refGene set.

Mouse genes in a custom non-redundant set (n = 20,633) (Rose et al. 2016) were classified into three groups based on the overlap of their gene promoters with non-methylated CpG islands (NMI), as well as RING1B- and SUZ12-bound sites. NMIs (n = 27,047) were identified using MACS2 peak calling with the matching input control from BioCAP-seq (Long et al. 2013). All genes with promoters (TSS ± 1 kb) not overlapping with NMIs were referred to as non-NMI genes (n = 6333). NMI-overlapping genes were further sub-divided into PcG-occupied genes (n = 5582), if their promoters overlapped with both RING1B- and SUZ12-bound sites defined in a previous study (Blackledge et al. 2020), and non-PcG-occupied genes (n = 8718), if they did not. The overlap between NMIs, RING1B- and SUZ12-bound regions with gene promoters (TSS ± 1 kb) was determined using the closest function from BEDTools.

### Read count quantitation and analysis

For cChIP-seq and cATAC-seq, computeMatrix and plotProfile/plotHeatmap from deeptools were used to perform metaplot and heatmap analysis of read density at regions of interest. Metaplot profiles represented the mean read density over a set of genomic regions, except for Pol II cChIP-seq, for which the median read density was plotted due to an extremely broad range of signal intensities across the intervals of interest. For chromosome-wide density plots, read coverage in 250 kb bins was calculated using a custom R script utilising GenomicRanges, GenomicAlignments and Rsamtools Bioconductor packages (Huber et al. 2015) and visualised using ggplot2. For cChIP-seq and cATAC-seq, target regions of interest were annotated with read counts from merged spike-in normalised replicates using multiBamSummary from deeptools (“—outRawCounts”). For differential gene expression analysis, we used a custom Perl script utilising SAMtools (Li et al. 2009) to obtain read counts from individual biological replicates prior to spike-in normalisation for a custom non-redundant mm10 gene set (n = 20,633).

Normalised read counts and log2-fold changes for different genomic intervals were visualised using custom R scripts and ggplot2. For boxplot analysis of cChIP-seq and cATAC-seq signal, read counts were normalised to the genomic region size (in kb). For boxplots, boxes show interquartile range (IQR) and whiskers extend by 1.5xIQR. ggcor function from the GGally R package was used to generate a correlation matrix for the association between log2-fold changes in gene expression (cnRNA-seq) and log2-fold changes in cChIP-seq signal for Pol II and transcription-associated histone modifications at gene promoters or bodies following OHT treatment in *Bap1*^*fl/fl*^ ESCs. All correlation analyses used Pearson correlation coefficient to measure the strength of the association between the variables and were visualised using scatterplots colored by density with stat_density2d. Linear regression was plotted using stat_poly_eq function from the ggpmisc R package, together with the model’s R^2^_adj_ coefficient of determination.

### Differential gene expression analysis

To identify significant gene expression changes in cnRNA-seq, we used a custom R script that incorporates spike-in calibration into DESeq2 analysis (Love et al. 2014) as described previously (Fursova et al. 2019; Blackledge et al. 2020). Briefly, dm6 read counts were obtained for unique dm6 refGene genes to calculate DESeq2 size factors for normalisation of raw mm10 read counts for a custom non-redundant mm10 gene set. Prior to quantification, dm6 reads were pre-normalised using the dm6/mm10 total read ratio in the corresponding gDNA-seq samples in order to account for variations in spike-in cell mixing. For visualisation and ranking of the effect sizes, we performed shrinking of log2-fold changes using the original DESeq2 shrinkage estimator with an adaptive normal distribution as prior (Love et al. 2014). For visualisation of DESeq2-normalized read counts, they were averaged across the replicates and used to calculate RPKM. For a change to be called significant, we applied a threshold of p-adj < 0.05 and fold change > 1.5. Log2-fold change values were visualised using R and ggplot2 with MA plots, heatmaps and boxplots/violinplots. ComplexHeatmap R package (Gu et al. 2016) was used to plot heatmaps of log2-fold gene expression changes. For MA plots, the density of the data points across y-axis was shown to reflect the general direction of gene expression changes. HOMER v4.9.1 (Heinz et al. 2010) was used to perform gene ontology (GO) term enrichment analysis for differentially expressed genes, with the custom non-redundant mm10 gene set used as background.

## Data and software availability

The high-throughput sequencing data reported in this study have been deposited in GEO under the accession number GSEXXXXXX. Published data used in this study include BioCAP-seq, GEO:GSE43512 (Long et al. 2013); cnRNA-seq in *PRC1*^*CKO*^ ESCs, GEO:GSE119619 (Fursova et al. 2019); RING1B- and SUZ12-bound regions in ESCs, GEO:GSE132752 (Blackledge et al. 2020); ATAC peaks from E14 ESCs, GEO:GSE98403 (King et al. 2018); CTCF ChIP-seq in wild-type ESCs, GEO:GSE153400 (Huang et al. 2020); ChIP-seq for OCT4, NANOG and SOX2 in untreated *Brg1*^*fl/fl*^ ESCs, GEO:GSE87822 (King and Klose 2017); H3K9me3 and H3K36me3 ChIP-seq in LIF-grown ESCs, GEO:GSE120376 (Ramisch et al. 2019). All R and Perl scripts used for data analysis in this study are available upon request.

---

**Supplemental Figure S1.**
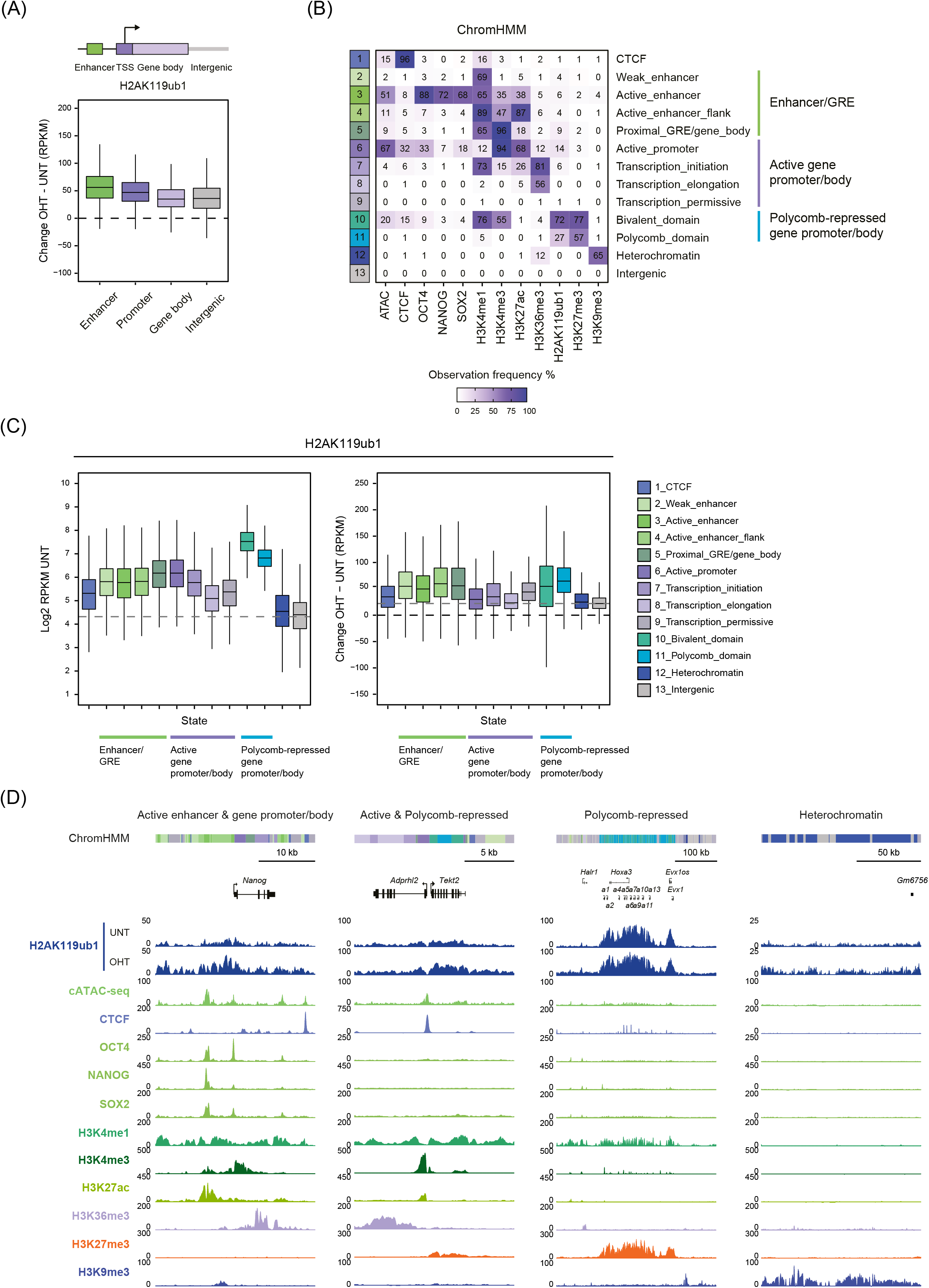
Related to Figure 1. **(A)** Boxplots showing the absolute change in H2AK119ub1 cChIP-seq signal at gene regulatory elements (enhancers and promoters), gene bodies, and intergenic regions following OHT treatment in *Bap1*^*fl/fl*^ ESCs. **(B)** A heatmap summarizing emission parameters for the ChromHMM model used to segment the genome into 13 chromatin states. Each row of the heatmap corresponds to one of the chromatin states, which are color-coded and grouped based on the underlying gene regulatory elements (GREs) and transcriptional activity. Columns correspond to different chromatin features that were used to build the model. The heatmap color intensity reflects the probability of observing a particular chromatin feature in the specific chromatin state. **(C)** Boxplots comparing cChIP-seq signal for H2AK119ub1 in untreated *Bap1*^*fl/fl*^ ESCs (*left panel*), as well as the absolute change in this signal following OHT treatment in *Bap1*^*fl/fl*^ ESCs *(right panel)*, across different chromatin states defined by the ChromHMM model in (B). The dashed grey line represents the average levels of H2AK119ub1 in untreated cells (*left panel*) or the average change in H2AK119ub1 following BAP1 removal *(right panel)* across the genome, as determined by their median values in intergenic regions. **(D)** Snapshots of genomic regions encompassing different chromatin states defined by ChromHMM. ChIP-seq tracks for all chromatin features that were used to build the ChromHMM model in (B) are shown together with H2AK119ub1 cChIP-seq in untreated and OHT-treated *Bap1*^*fl/fl*^ ESCs. The segmentation of genomic regions into chromatin states is illustrated with the colored bar at the top of each panel, with different states represented by the same colors as in (B).

**Supplemental Figure S2.**
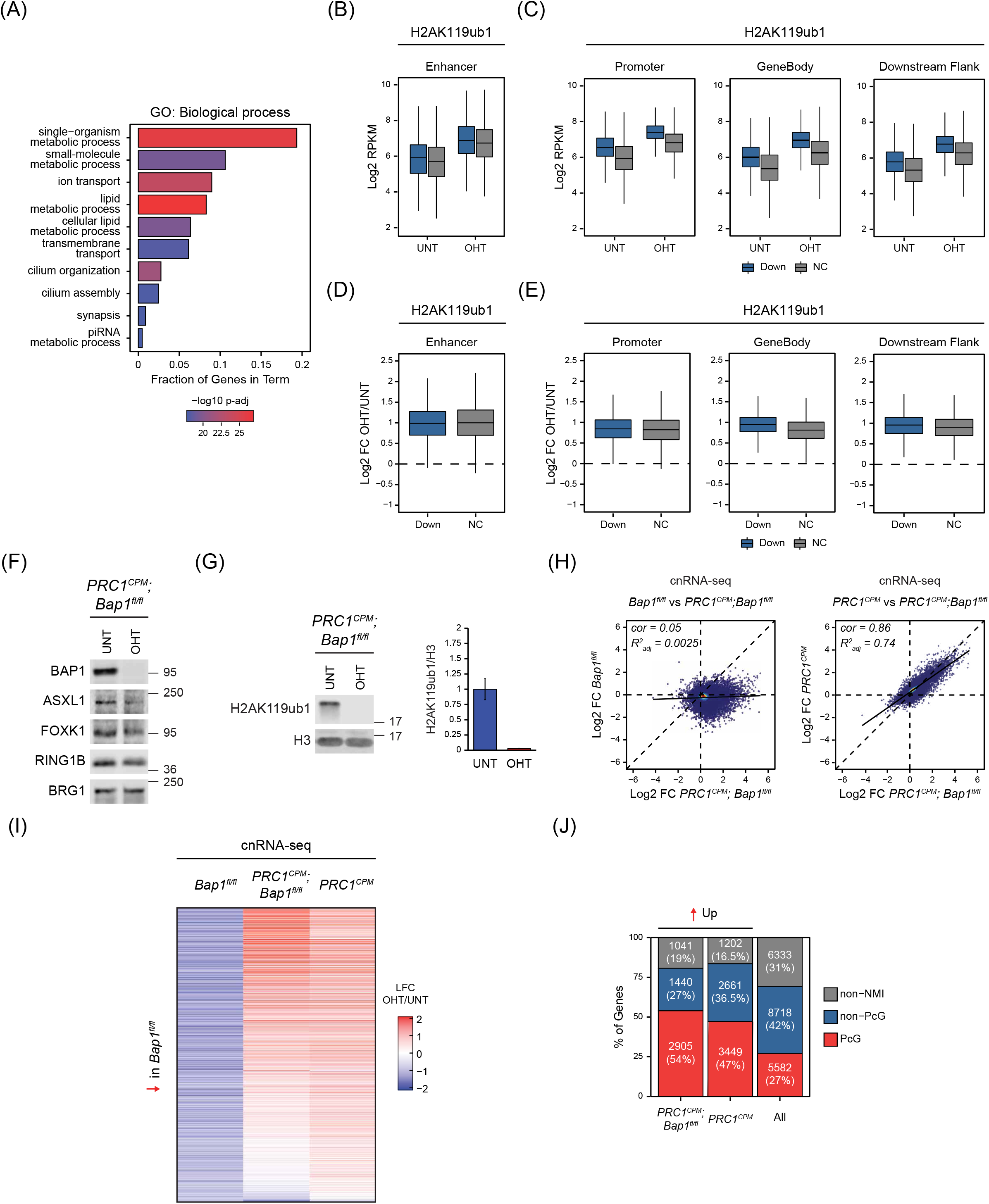
Related to Figure 2. **(A)** A gene ontology (GO) analysis of Biological Process term enrichment for genes showing a significant reduction in expression (p-adj < 0.05 and > 1.5-fold) in *Bap1*^*fl/fl*^ cells following OHT treatment. **(B)** Boxplots comparing H2AK119ub1 cChIP-seq signal in *Bap1*^*fl/fl*^ ESCs (untreated and OHT-treated) at the nearest putative enhancers associated with genes that show a significant reduction (Down, n = 1916) or no change (No Change, n = 9647) in expression following BAP1 removal based on cnRNA-seq analysis (p-adj < 0.05 and > 1.5-fold). **(C)** Boxplots comparing H2AK119ub1 cChIP-seq signal in *Bap1*^*fl/fl*^ ESCs (untreated and OHT-treated) at the promoters, bodies and 10 kb downstream flanking regions of genes that show a significant reduction (Down, n = 2828) or no change (No Change, n = 17,203) in expression following BAP1 removal based on cnRNA-seq analysis (p-adj < 0.05 and > 1.5-fold). **(D)** Boxplots comparing log2-fold changes in H2AK119ub1 cChIP-seq signal following OHT treatment in *Bap1*^*fl/fl*^ ESCs at the nearest putative enhancers associated with genes that were split into two groups as defined in (B). **(E)** Boxplots comparing log2-fold changes in H2AK119ub1 cChIP-seq signal following OHT treatment in *Bap1*^*fl/fl*^ ESCs at the promoters, bodies, and 10 kb downstream flanking regions of genes that were split into two groups as defined in (C). **(F)** Western blot analysis for the PR-DUB complex subunits (BAP1, ASXL1 and FOXK1) and the PRC1 catalytic subunit RING1B in untreated and OHT-treated *PRC1*^*CPM*^;*Bap1*^*fl/fl*^ ESCs. BRG1 is shown as a loading control. **(G)** Western blot analysis (*left panel*) and quantification (*right panel*) of H2AK119ub1 levels relative to histone H3 in untreated and OHT-treated *PRC1*^*CPM*^;*Bap1*^*fl/fl*^ ESCs. Error bars represent SEM (n = 3). **(H)** Scatterplots comparing the log2-fold changes in gene expression (cnRNA-seq) following OHT treatment in *Bap1*^*fl/fl*^ and *PRC1*^*CPM*^;*Bap1*^*fl/fl*^ ESCs (*left panel*), as well as *PRC1*^*CPM*^ and *PRC1*^*CPM*^;*Bap1*^*fl/fl*^ ESCs (*right panel*). *R*^*2*^_*adj*_ represents the adjusted coefficient of determination for linear regression, and *cor* denotes the Pearson correlation coefficient. This illustrates that simultaneous removal of BAP1 and catalytic activity of PRC1 closely recapitulates the gene expression defects manifesting from disrupting PRC1 catalysis alone, revealing an epistatic genetic interaction between PRC1 (H2AK119ub1) and BAP1. **(I)** A heatmap illustrating log2-fold changes in expression (cnRNA-seq) following OHT treatment (LFC OHT/UNT) in *Bap1*^*fl/fl*^, *PRC1*^*CPM*^;*Bap1*^*fl/fl*^ and *PRC1*^*CPM*^ ESCs for genes that show a significant reduction in expression (p-adj < 0.05 and > 1.5-fold) after BAP1 removal. **(J)** A bar plot illustrating the distribution of different gene classes among genes that become significantly derepressed (p-adj < 0.05 and > 1.5-fold) following OHT treatment in *PRC1*^*CPM*^;*Bap1*^*fl/fl*^ and *PRC1*^*CPM*^ ESCs. PcG corresponds to Polycomb-occupied genes; Non-PcG to non-Polycomb-occupied genes; Non-NMI to genes lacking a non-methylated CGI (NMI) at their promoter.

**Supplemental Figure S3.**
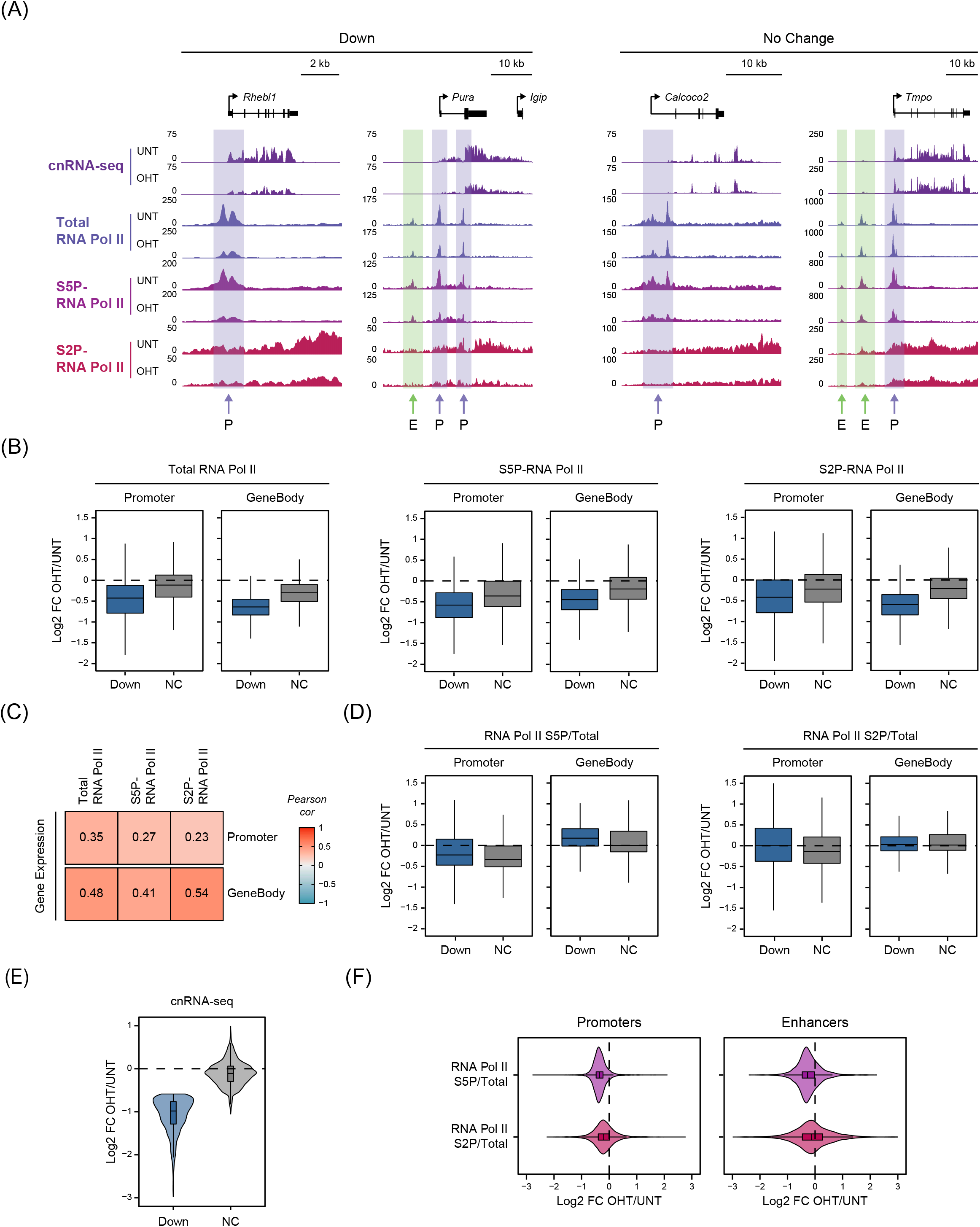
Related to Figure 3. **(A)** Snapshots of genes that display a significant reduction (Down) or no change in expression following removal of BAP1 based on cnRNA-seq analysis (p-adj < 0.05 and > 1.5-fold). Gene expression (cnRNA-seq) and cChIP-seq for total Pol II, as well as its Ser5- and Ser2-phosphorylation (S5P- and S2P-Pol II), are shown in *Bap1*^*fl/fl*^ ESCs (untreated and OHT-treated). Positions of promoters (P) (H3K27ac-high, H3K4me3-high) and the nearest putative enhancers (E) (H3K27ac-high, H3K4me3-low) for these genes are indicated (also see Figure S4A). **(B)** Boxplots comparing log2-fold changes in cChIP-seq signal for total Pol II, as well as its Ser5- and Ser2-phosphorylation (S5P- and S2P-Pol II), following OHT treatment in *Bap1*^*fl/fl*^ ESCs at promoters and over the bodies of genes that show a significant reduction (Down, n = 2828) or no change (No Change, n = 17,203) in expression after BAP1 removal based on cnRNA-seq analysis (p-adj < 0.05 and > 1.5-fold). **(C)** Correlation of log2-fold changes in gene expression (cnRNA-seq) with log2-fold changes in cChIP-seq signal for total Pol II, as well as its Ser5- and Ser2-phosphorylation (S5P- and S2P-Pol II), at gene promoters and bodies in *Bap1*^*fl/fl*^ cells following OHT treatment. **(D)** Boxplots comparing log2-fold changes in the abundance of Ser5- and Ser2-phosphorylation relative to total Pol II levels (S5P/Total and S2P/Total) following OHT treatment in *Bap1*^*fl/fl*^ ESCs at promoters and over the bodies of genes that were split into two groups as defined in (B). **(E)** Violinplots comparing log2-fold changes in gene expression (cnRNA-seq) following OHT treatment in *Bap1*^*fl/fl*^ ESCs for genes that show a significant reduction (Down, n = 2828) or no change (No Change, n = 17,203) in expression after BAP1 removal based on cnRNA-seq analysis (p-adj < 0.05 and > 1.5-fold). This illustrates that expression of genes that are not classified as showing significant changes is still reduced following BAP1 removal, albeit very modestly. **(F)** Violinplots comparing log2-fold changes in the abundance of Ser5- and Ser2-phosphorylation relative to total Pol II levels (S5P/Total and S2P/Total) following OHT treatment in *Bap1*^*fl/fl*^ ESCs at active promoters and enhancers.

**Supplemental Figure S4.**
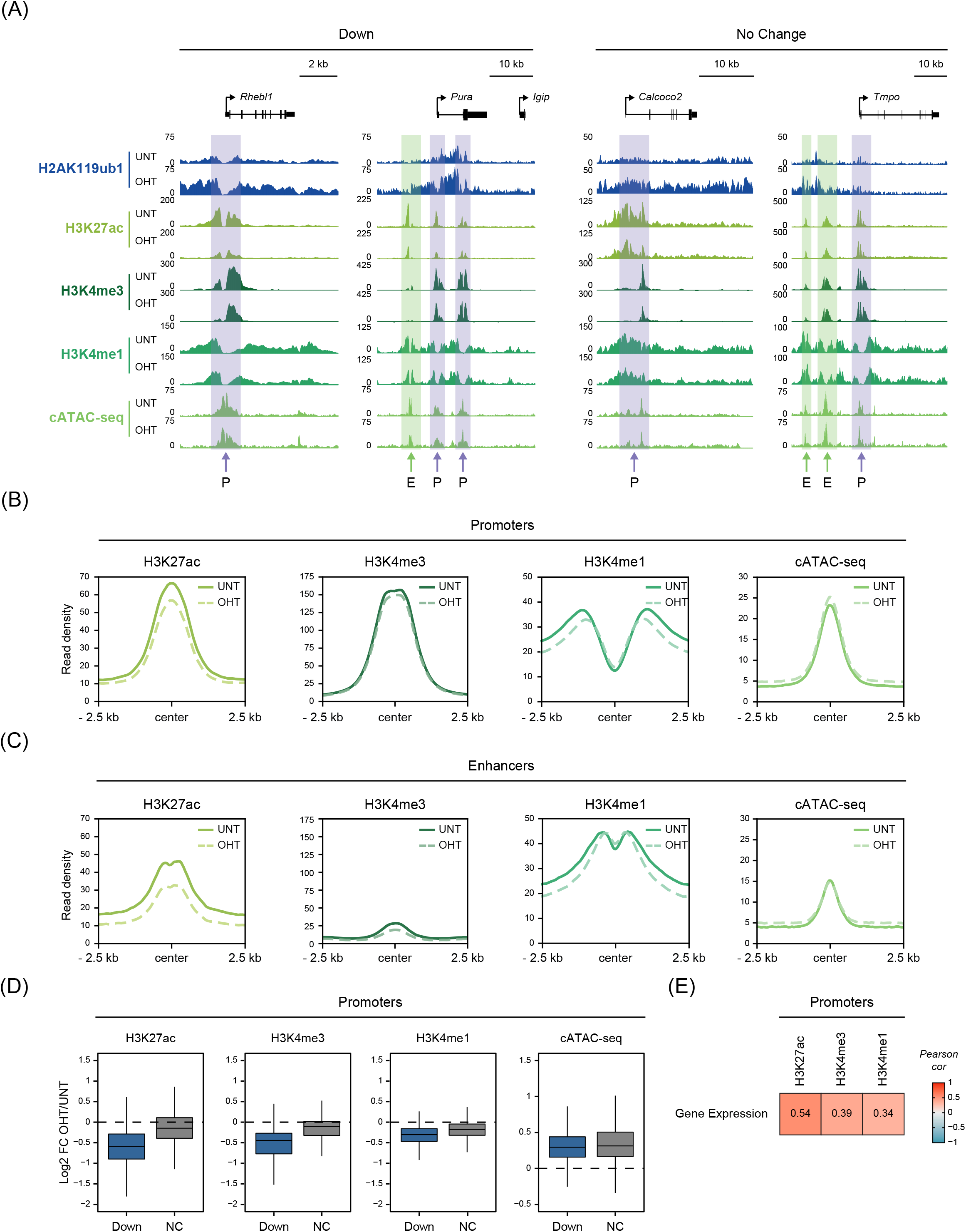
Related to Figure 4. **(A)** Snapshots of genes that display a significant reduction (Down) or no change in expression following removal of BAP1 based on cnRNA-seq analysis (p-adj < 0.05 and > 1.5-fold). cChIP-seq is shown for H2AK119ub1, H3K27ac, H3K4me3 and H3K4me1 in *Bap1*^*fl/fl*^ ESCs (untreated and OHT-treated). cATAC-seq is also shown as a measure of chromatin accessibility. Positions of promoters (P) (H3K27ac-high, H3K4me3-high) and the nearest putative enhancers (E) (H3K27ac-high, H3K4me3-low) for these genes are indicated. **(B)** Metaplots illustrating H3K27ac, H3K4me3 and H3K4me1 cChIP-seq signal, as well as cATAC-seq signal, at active gene promoters in *Bap1*^*fl/fl*^ ESCs (untreated and OHT-treated). **(C)** As in (B) but for active enhancers. **(D)** Boxplots comparing log2-fold changes in H3K27ac, H3K4me3 and H3K4me1 cChIP-seq signal, as well as in cATAC-seq signal, following OHT treatment in *Bap1*^*fl/fl*^ ESCs at the promoters of genes that show a significant reduction (Down, n = 2828) or no change (No Change, n = 17,203) in expression after BAP1 removal based on cnRNA-seq analysis (p-adj < 0.05 and > 1.5-fold). **(E)** Correlation of log2-fold changes in gene expression (cnRNA-seq) with log2-fold changes in cChIP-seq signal for H3K27ac, H3K4me3 and H3K4me1 at gene promoters following OHT treatment in *Bap1*^*fl/fl*^ cells.

**Supplemental Figure S5.**
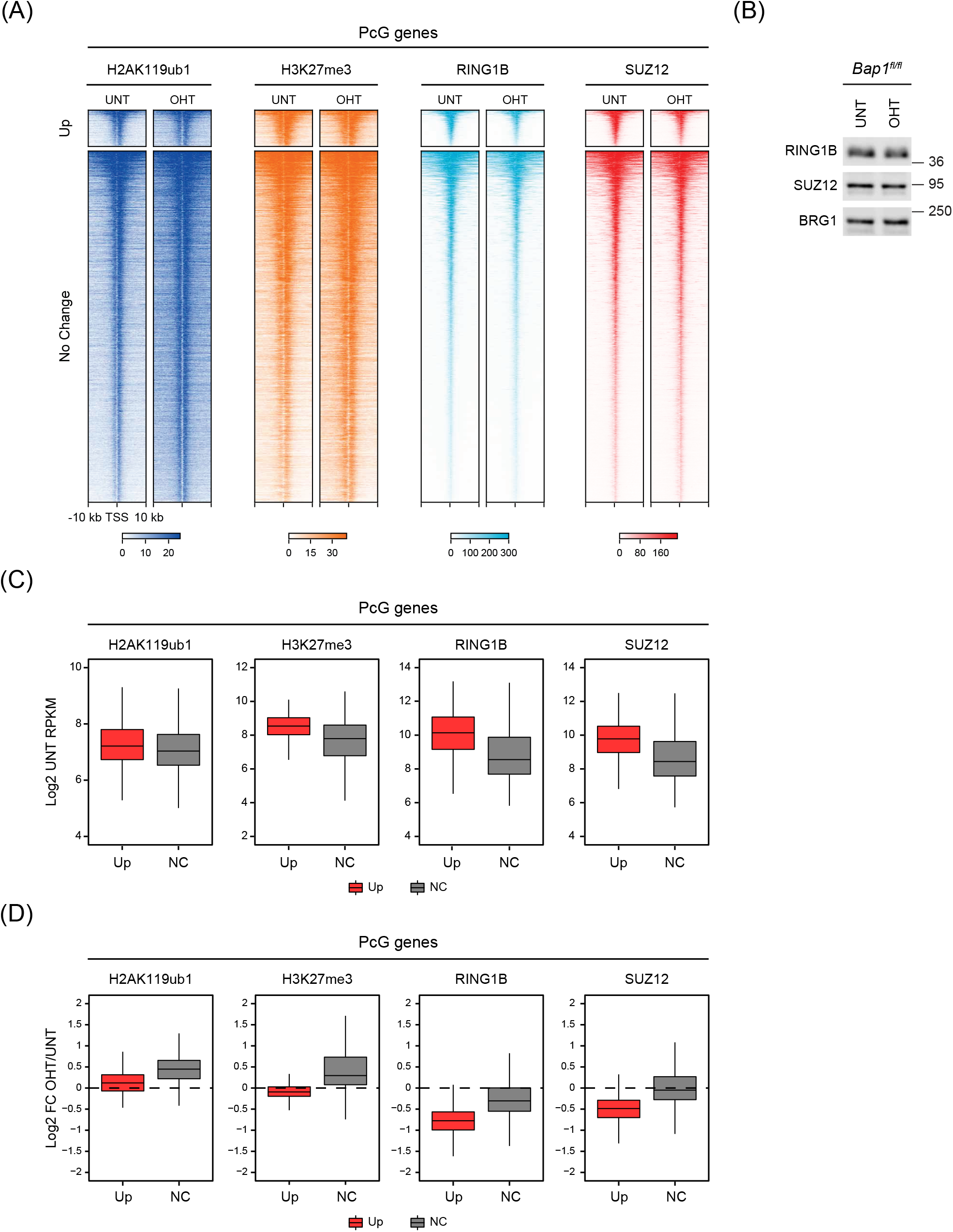
Related to Figure 5. **(A)** Heatmaps of cChIP-seq signal for H2AK119ub1, H3K27me3, RING1B (PRC1) and SUZ12 (PRC2) in *Bap1*^*fl/fl*^ ESCs (untreated and OHT-treated) at the promoters of Polycomb target genes that become significantly derepressed (Up, n = 421) or do not change in expression (No Change, n = 4075) following BAP1 removal based on cnRNA-seq analysis (p-adj < 0.05 and > 1.5-fold). Intervals were sorted by RING1B occupancy in untreated *Bap1*^*fl/fl*^ ESCs. **(B)** Western blot analysis for RING1B (PRC1) and SUZ12 (PRC2) in untreated and OHT-treated *Bap1*^*fl/fl*^ ESCs. BRG1 is shown as a loading control. **(C)** Boxplots comparing cChIP-seq signal for H2AK119ub1, H3K27me3, RING1B and SUZ12 in untreated *Bap1*^*fl/fl*^ ESCs at the promoters of Polycomb target genes that become significantly derepressed (Up, n = 421) or do not change in expression (No Change, n = 4075) following BAP1 removal based on cnRNA-seq analysis (p-adj < 0.05 and > 1.5-fold). **(D)** Boxplots comparing log2-fold changes in cChIP-seq signal for H2AK119ub1, H3K27me3, RING1B and SUZ12 following OHT treatment in *Bap1*^*fl/fl*^ ESCs at the promoters of Polycomb target genes that become significantly derepressed (Up, n = 421) or do not change in expression (No Change, n = 4075) after BAP1 removal.

